# Integrative deep immune profiling of the elderly reveals systems-level signatures of aging, sex, smoking, and clinical traits

**DOI:** 10.1101/2024.07.10.602828

**Authors:** Lennart Riemann, Rodrigo Gutierrez, Ivan Odak, Joana Barros-Martins, Lennart M. Roesner, Ximena Leon Lara, Christine Falk, Thomas F. Schulz, Gesine Hansen, Thomas Werfel, Reinhold Förster, the RESIST SI Cohort Investigators

## Abstract

Elderly individuals have higher disease susceptibility and lower vaccine responsiveness, highlighting the need to better comprehend the aging immune system and its clinical associations. Here we conducted a deep immune profiling study of 550 elderly individuals (61–94 years) and 100 young adults (22–38 years). Utilizing high-dimensional spectral flow cytometry to identify 97 immune cell populations and 48-plex cytokine profiling, we detailed intricate age-and sex-related changes in the elderly immune system at an unprecedented depth. Synthesizing information from clinical, laboratory, and immunological data through an integrative multi-block analysis, we reveal overarching systems-level signatures of aging, such as increased concentrations of specific cytokines and frequencies of defined innate and adaptive immune cell subpopulations. Extending this approach, we identified unique immune signatures of smoking, obesity, and several diseases including osteoporosis, heart failure and gout. Our systems biology approach enables to uncover new relationships between clinical characteristics and immunological traits.

## Main

The global population is aging at an unprecedented pace^1^ and more than one in six people worldwide will be older than 60 years by 2030.^2^ The incidence, prevalence, and severity of many infectious and non-infectious diseases increase with age, while vaccine responses diminish, rendering disease prevention more difficult.^3–6^ As a result, understanding the complexities of the immune system in relation to aging, age-related diseases, and other patient characteristics, such as smoking and obesity, is becoming increasingly important and could be an avenue to find new therapies and interventions.

Understanding the complexities of the immune system, however, remains challenging due to the vast array of interacting immune parameters, including hundreds of different cell types and signaling molecules, and their high inter-individual variability in humans.^7,8^ The arrival and improvement of new technologies, such as mass and spectral flow cytometry, multiplex assays, and high-throughput sequencing, has dramatically increased the number of parameters that can be analyzed simultaneously, enabling to study the human immune system at a far higher resolution and in a more comprehensive manner than ever before. However, the high-dimensionality of the acquired data necessitates novel computational approaches for efficient yet biologically informative analysis, and large sample sizes to detect even subtle changes efficiently in highly variable biological systems. Such large-scale studies have significantly enhanced our understanding of the human immune system in various contexts of health and disease.^9–16^

Here we report a comprehensive, deep immune phenotyping study of the aging human immune system and several age-related diseases in a large, well-characterized cohort of elderly individuals who had been randomly recruited from the general population. Our analysis revealed significant changes in cytokine/chemokine profiles and the composition of the innate and adaptive immune cell compartments across age and sex. Additionally, we fine-map previously reported associations, e.g., with smoking and cytomegalovirus status, to precise cell subpopulations and identify several novel associations with various diseases. Finally, using a innovative multi-block data fusion and integration approach to jointly analyze clinical, laboratory, and immunological data, we revealed integrative signatures of aging and of several age-related diseases across datasets, generating important new insights on a systems biology level. Collectively, our high-resolution immune profiling significantly advances our understanding of immune signatures of aging, sex, and other pertinent patient characteristics, such as smoking, obesity, and clinical diseases.

## The RESIST Senior Individuals Cohort

The RESIST Senior Individuals (SI) Cohort consists of 550 elderly individuals aged 61–94 years and 100 young adults aged 22–38 years. Participants were randomly selected from the general population through the local residents’ registry office and invited to participate in the study.^17^ Upon obtaining consent, participants underwent a comprehensive clinical questionnaire, physical exam, laboratory analyses, and collection of various biomaterials (**Fig. 1A**). Details on the study design and participant demographics have been described before.^17^

**Fig. 1:**
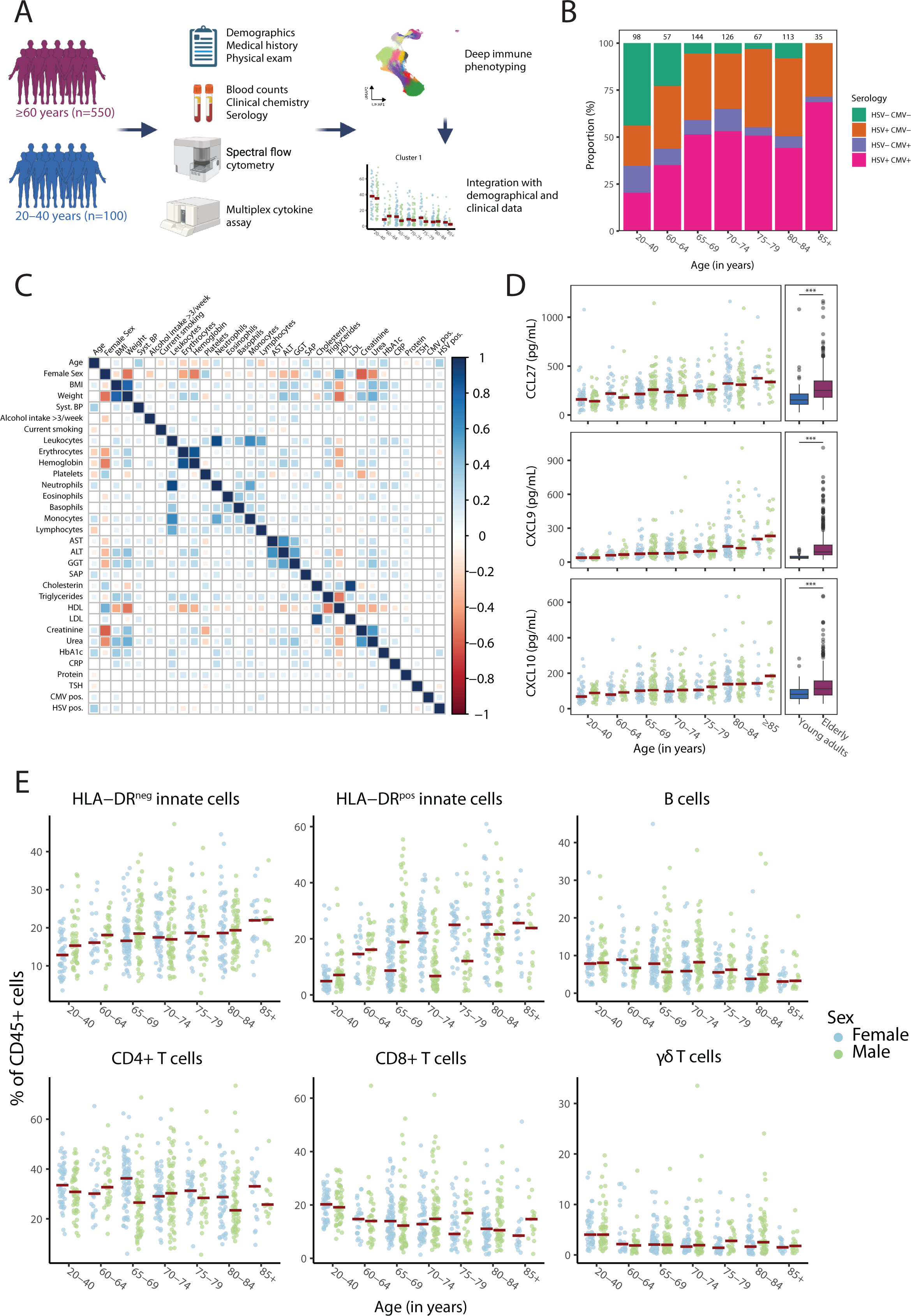
Studying the aging immune system in a large cohort of elderly individuals. (**A**) Study overview. (**B**) Proportion of HSV and CMV seropositive study participants across age groups. Numbers on top of the bars indicate the number of individuals within each age group. (**C**) Heatmap displaying Spearman correlation coefficients between indicated clinical and laboratory features in the elderly participants. Only significant correlations (p ≤0.05) are shown. (**D**) Scatter plots showing the plasma concentrations of CCL27, CXCL9, and CXCL10 in females (blue) and males (green) across age groups. The red crossbar indicates the median in each group. Plasma levels of these cytokines were significantly higher in elderly compared to young adults (Wilcoxon rank sum test, p < 0.001). (**E**) Frequencies of pre-gated major immune cell subsets in females (blue) and males (green) across age groups. The red crossbar indicates the median in each group.

Baseline characteristics of the elderly individuals including comorbidities reflected a typical aging Western society and are summarized in **Table 1**. The sex distribution was balanced between males and females. Thirty-five percent of young adults were serologically positive for CMV, while this proportion increased to 58% in elderly individuals (**Fig. 1B**). Similarly, 41% and 85% of young adults and elderly, respectively, were herpes simplex virus (type 1 or 2, HSV) positive, and almost all participants were seropositive for varicella zoster virus (**Table 1**). Significant correlations cross-linking demographical, anthropometrical, laboratory and serological data are presented in **Fig. 1C**, revealing known associations such as increasing creatinine levels and systolic blood pressure with age. Strong correlations were also found among different circulating immune cells, as well as among liver enzymes, and plasma lipids. Body mass index (BMI) correlated significantly with HbA1c levels, as expected, but interestingly, a significant positive correlation was also observed between HbA1c levels and HSV seropositivity, in line with a recent population-based study reporting a possible link between HSV type 2 infection and prediabetes.^18^ In a multiplex analysis of 48 cytokines, we found plasma concentrations of several pro-inflammatory chemokines, such as CCL27, CXCL9, and CXCL10, to increase with age (**Fig. 1D**).This indicates a propensity towards a state of low-grade inflammation, sometimes termed inflammaging^19^, a key feature of the aging immune system. While CCL3 (MIP1α), G-CSF, and HGF were also elevated in the elderly, notably, other classical pro-inflammatory cytokines commonly associated with inflammaging, including TNF-α, interferon-γ, and IL-17, did not exhibit statistically significant differences between young adults and elderly participants (**Suppl. Table 1**).

**Table 1:** Demographical, anthropometrical, clinical, and serological characteristics of participants in the RESIST SI cohort.

| Variable | Young adults<br>(N=100) | Elderly<br>(N=550) | P value | Missing data |  |
| --- | --- | --- | --- | --- | --- |
|  |  |  |  | Young<br>adults | Elderly |
| Age (years) |  |  |  | 0 (0%) | 0 (0%) |
| Median | 29 (25–32) | 73 (68–80) | <0.001 |  |  |
| ≥70 years | 0 (0%) | 347 (63%) | <0.001 |  |  |
| ≥80 years | 0 (0%) | 148 (27%) | <0.001 |  |  |
| Sex |  |  | 0.744 | 0 (0%) | 0 (0%) |
| Female (%) | 48 (48%) | 277 (50%) |  |  |  |
| Male (%) | 52 (52%) | 273 (50%) |  |  |  |
| Weight (kg) | 68 (62–83) | 75 (65–85) | 0.026 | 1 (1%) | 9 (2%) |
| BMI (kg/m<sup>2</sup>) |  |  |  | 1 (1%) | 10 (2%) |
| Median | 24 (21–25) | 26 (23–29) | <0.001 |  |  |
| ≥25 (overweight) | 29 (29%) | 316 (59%) | <0.001 |  |  |
| ≥30 (obese) | 6 (6%) | 125 (23%) | <0.001 |  |  |
| Blood pressure (mmHg) |  |  |  | 1 (1%) | 5 (<1%) |
| Systolic | 119 (112–130) | 136 (126–148) | <0.001 |  |  |
| Diastolic | 75 (70–80) | 78 (71–85) | 0.001 |  |  |
| Syst. ≥140 mmHg | 11 (11%) | 235 (43%) | <0.001 |  |  |
| Alcohol |  |  | <0.001 | 1 (1%) | 9 (2%) |
| Never | 12 (12%) | 70 (13%) |  |  |  |
| <1/month | 22 (22%) | 98 (18%) |  |  |  |
| 2–4/month | 47 (48%) | 136 (25%) |  |  |  |
| 2–3/week | 14 (14%) | 128 (24%) |  |  |  |
| ≥4/week | 4 (4%) | 109 (20%) |  |  |  |
| Smoking |  |  |  |  |  |
| Ever | 26 (26%) | 296 (54%) | <0.001 | 1 (1%) | 3 (<1%) |
| Current | 15 (16%) | 55 (10%) | 0.138 | 6 (6%) | 8 (1%) |
| Medical history |  |  |  |  |  |
| Hypertension | 6 (6%) | 310 (57%) | <0.001 | 0 (0%) | 1 (<1%) |
| Myocardial infarction | 0 (0%) | 30 (5%) | 0.009 | 0 (0%) | 1 (<1%) |
| Heart failure | 1 (1%) | 40 (7%) | 0.031 | 0 (0%) | 1 (<1%) |
| Cardiac arrhythmias | 2 (2%) | 118 (22%) | <0.001 | 0 (0%) | 1 (<1%) |
| Cancer | 1 (1%) | 137 (25%) | <0.001 | 0 (0%) | 3 (<1%) |
| Diabetes mellitus | 1 (1%) | 53 (10%) | 0.007 | 0 (0%) | 2 (<1%) |
| Thyroid disease | 10 (10%) | 130 (24%) | <0.001 | 0 (0%) | 2 (<1%) |
| COPD | 2 (2%) | 41 (7%) | 0.074 | 1 (1%) | 1 (<1%) |
| Allergies | 45 (46%) | 229 (42%) | 0.515 | 2 (2%) | 2 (<1%) |
| Shingles | 9 (9%) | 100 (18%) | 0.036 | 1 (1%) | 2 (<1%) |
| Osteoporosis | 0 (0%) | 60 (11%) | 0.001 | 0 (0%) | 1 (<1%) |
| Gout | 0 (0%) | 35 (6%) | 0.018 | 0 (0%) | 2 (<1%) |
| Appendectomy | 8 (8%) | 190 (35%) | <0.001 | 0 (0%) | 0 (0%) |
| Tonsillectomy | 12 (12%) | 222 (40%) | <0.001 | 0 (0%) | 0 (0%) |
| Depression | 7 (7%) | 58 (11%) | 0.417 | 5 (5%) | 9 (2%) |
| Serology |  |  |  |  |  |
| HSV IgG positive | 41 (41%) | 466 (85%) | <0.001 | 1 (1%) | 3 (<1%) |
| VZV IgG positive | 99 (100%) | 544 (100%) | 1.000 | 1 (1%) | 3 (<1%) |
| CMV IgG positive | 35 (35%) | 314 (58%) | <0.001 | 1 (1%) | 4 (<1%) |

Numbers and percentages are displayed for categorical variables, while the median and interquartile range is displayed for continuous variables. Abb.: BMI: Body Mass Index; COPD: Chronic Obstructive Pulmonary Disease; CMV: Cytomegalovirus; HSV: Herpes Simplex Virus (type 1 or 2); VZV: Varicella Zoster Virus.

We then conducted an in-depth pan-analysis of the leukocyte composition of the elderly immune system by employing spectral flow cytometry on collected peripheral blood mononuclear cells. To achieve high-resolution phenotyping of both the innate and adaptive immune cell compartments, samples were first manually pre-gated on major immune cell lineages (HLA-DR^neg^ innate cells, HRA-DR^pos^ innate cells, γδ T cells, B cells, CD4+ T cells, and CD8+ T cells; **Extended Data Fig. 1**) before an unsupervised clustering algorithm was utilized on all 650 samples to assign pre-gated cells to specific cluster subpopulations characterized by distinct global marker expression profiles. As depicted in **Fig. 1E**, the proportions of major immune cell populations exhibited substantial changes with age. The proportion of myeloid cells among all leukocytes was increasing. Conversely, the proportions of B cells, CD4+ T cells, CD8+ T cells, and γδ T cells decreased with age. Subsequent unsupervised clustering of cells within these pre-gated populations allowed us to identify a total of 97 cell subpopulations with distinct immunophenotypic profiles, including 36 innate and 61 adaptive immune cell types. We then studied their characteristics and age-related trends in detail, and integrated their frequencies with available clinical and serological data.

## Innate immune cell compartment

We first investigated HLA-DR^neg^ innate cells, which were clustered into 15 distinct clusters consisting of different types of natural killer cells (NK cells; clusters 1–8), basophils (cluster 9), mast cells (cluster 10), innate lymphoid cells (ILC; clusters 11–14), and one small cluster (cluster 15) that could not be assigned to a specific cell population. Marker characteristics of all clusters are depicted in **Fig. 2A** and **Extended Data Fig. 2A**. While the two largest clusters (clusters 4 and 5, mature NK) remained fairly stable across age, notable changes were seen in basophils and mast cells, which slightly increased with age, and in ILCs type 2 and 3, which contracted with age (**Fig. 2B**; **Extended Data Fig. 3**; **Suppl. Table 2**). Notably, seven of the 15 clusters were significantly associated with sex, particularly early and mature NK cell clusters 1–6 (**Fig. 2C**). Clusters 1, 2, and 4 tended to be higher in males, whereas clusters 3, 5, 6, and 15 were associated with female sex. Regarding potential associations with diseases and smoking, cluster 15 (ILC type 1) tended to be higher in smokers, while cluster 2 (early NK) was negatively associated with smoking. The large clusters 4 and 5 (mature NK cells) were significantly associated with obesity (BMI ≥30 kg/m^2^), but notably in opposing directions (**Fig. 2C**).

**Fig. 2:**
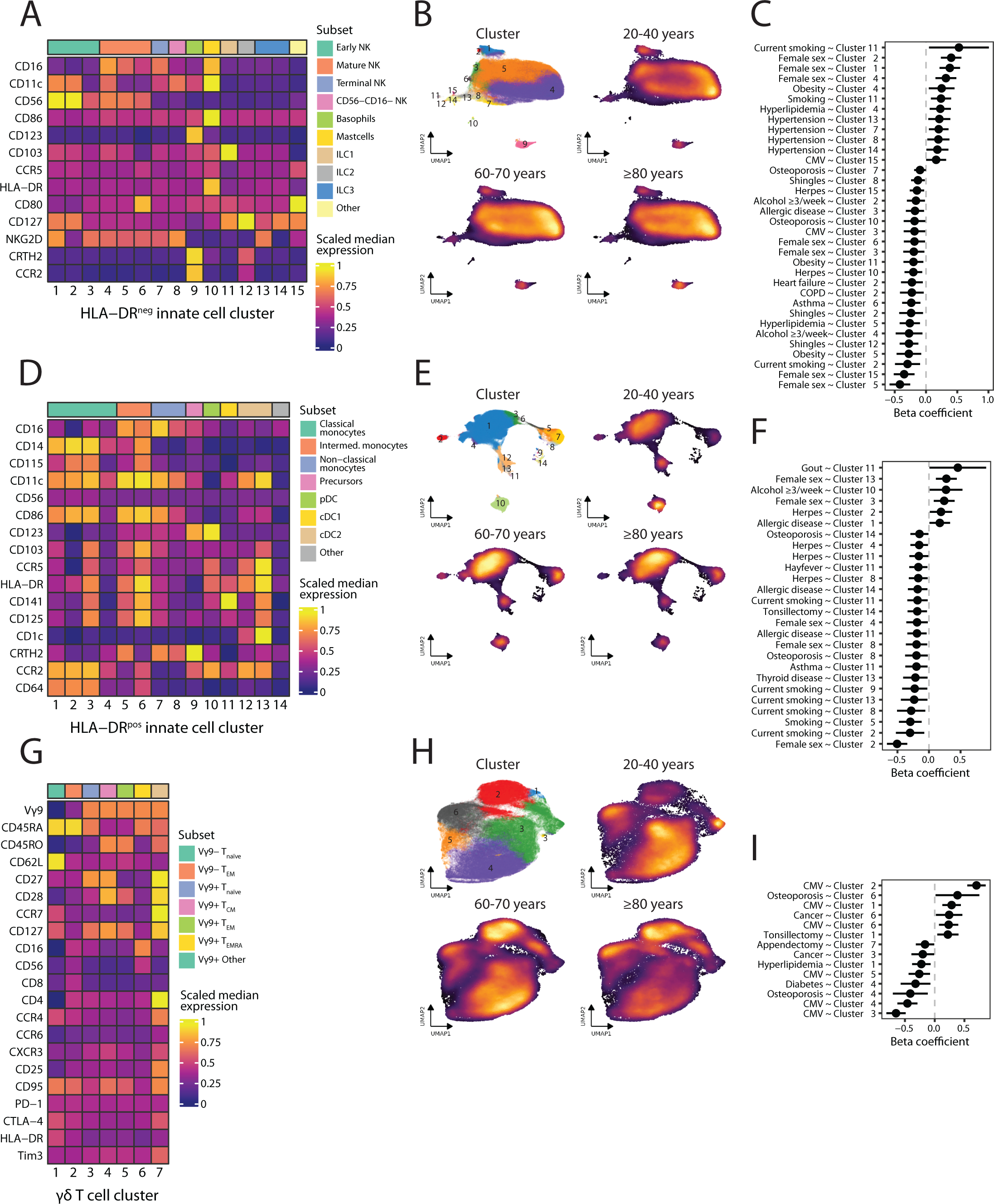
Marker characteristics, age-related changes, and associations of innate immune cells with clinical data. (**A, D, F**) Heatmap displaying transformed and row-scaled median marker expressions for each cluster of pre-gated HLA-DR^neg^ innate cells (**A**), HLA-DR^pos^ innate cells (**D**), and γδ T cells (**G**). Clusters were grouped into sub-lineage subset types based on canonical marker expressions (see methods). (**B**, **E**, **H**) UMAP cluster and density plots depicting the clusters and age distributions of HLA-DR^neg^ innate cells (**B**), HLA-DR^pos^ innate cells (**E**), and γδ T cells (**H**). (**C**, **F**, **I)**, Effect size plots with 0.95 confidence intervals from robust linear regression models with scaled cluster frequencies of HLA-DR^neg^ innate cells (**C**), HLA-DR^pos^ innate cells (**F**), and γδ T cells (**H**) as responses, and indicated variables as predictors. All models were adjusted for age and sex, except for models featuring sex, which were only adjusted for age. Only significant predictors (p ≤0.05) are displayed. Abbr.: Allergic disease: presence of at least one allergic disease, CMV: CMV seropositivity, Herpes: history of herpes symptoms (genital or labial), Shingles: history of shingles, Smoking: history of smoking, Thyroid disease: history of any thyroid disease.

Next, we studied HLA-DR^pos^ innate cells comprising 14 clusters of classical monocytes (clusters 1–3), intermediate monocytes (clusters 4–6), non-classical monocytes (clusters 7–8), granulo-monocytic precursors (cluster 9), peripheral dendritic cells (pDC; cluster 10), conventional dendritic cells (cDC) type 1 and 2 (clusters 11–13), and one cluster of cells negative for all measured markers (cluster 14; **Fig. 2D** and **Extended Data Fig. 2B**). Cluster 1 (classical monocyte), the largest HLA-DR^pos^ cluster, and cluster 4 (intermediate monocyte) remained largely stable across age groups, though with a high inter-individual variability. The frequencies of several monocyte clusters (clusters 3, 5, 6, 7, and 8) increased with age, while those of clusters 10 (pDC) and 12 (cDC2) decreased (**Fig. 2E**; **Extended Data Fig. 4**; **Suppl. Table 3**). One small monocyte cluster (cluster 3), distinctively expressing the tissue-residency marker CD103, increased with age and has not been described in the literature yet in the context of aging. Three clusters were significantly associated with sex, namely two classical monocyte clusters (clusters 2 and 3, in opposing directions), and cDC type 2 cluster 13, which tended to be higher in females (**Fig. 2F**). Cluster 2 (classical monocyte) was negatively associated with current smoking, consistent with a study from the UKTwin cohort^20^ describing decreased frequencies of classical monocyte subsets in active smokers (**Fig. 2F**). Cluster 11 (cDC type 1) was positively associated with gout, though the confidence interval was wide.

As part of the innate compartment, we identified 7 γδ T cell clusters encompassing naïve (cluster 1) and effector memory (T_EM_) Vγ9-T cells, naïve (cluster 3), central memory (T_CM_; cluster 4) and T_EM_ (cluster 5) Vγ9+ T cells, terminally differentiated (T_EMRA_; cluster 6) Vγ9+ T cells, as well as one small additional cluster with naïve features plus CD4 co-expression (cluster 7; **Fig. 2G**, **Extended Data Fig. 2C**). UMAP density plots suggested marked age-associated changes (**Fig. 2H**). The strongest changes affected the Vγ9-EM cluster 2, which increased with age, and the resting Vγ9+ cluster 3, which contracted (**Extended Data Fig. 5**, **Suppl. Table 4**), consistent with previous reports.^21–23^ In contrast to previous observations^23^, we did not observe significant sex differences in any of the γδ T cell clusters. Strong associations were found with CMV seropositivity (**Fig. 2I**). While previous studies have shown increased γδ T cells in CMV positive individuals^22,24^, we could further delineate these associations to specific clusters. While both Vγ9-clusters (clusters 1 and 2) as well as cluster 6 (Vγ9+ T_EMRA_) were positively associated with CMV status, Vγ9+ clusters 3– 5, most prominently the naïve cluster 3, displayed negative associations. Additionally, two Vγ9+ clusters (clusters 4 and 6) displayed significant, yet opposing, associations with a history of osteoporosis.

Altogether, our deep immune phenotyping approach revealed significant associations related to age and disease, alongside discernable sex differences in the innate immune cell compartment. Associations were often subtle and subject to a large variability, highlighting the need for a large-scale study to detect them.

## Adaptive immune cell compartment

The B cell compartment consisted of 18 clusters comprising naïve B cells (clusters 1–2), non-switched memory (clusters 3–5) and switched memory (clusters 6–11) B cells, transitional B cells (clusters 12–15), plasmablasts (clusters 16–17), and double-negative B cells (cluster 18). Marker characteristics are provided in **Fig. 3A** and **Extended Data Fig. 6A**. In line with a recent single-cell sequencing study^14^ observing overall stable B cell cluster frequencies across life, we detected only minor changes with regard to age (**Fig. 3B**; **Extended Data Fig. 7**). In some clusters, such as clusters 1 and 10, small differences could be seen in participants above 80 years. Cluster 14 (transitional B cell) was rarer in elderly individuals compared to young adults, but then remained rather stable within the elderly. While frequency changes were limited, we noticed that the inter-individual variability of all 18 cluster frequencies was consistently higher in elderly individuals compared to young adults (**Suppl. Table 5**). Cluster 1 (naïve B) was positively associated with female sex, while clusters 8 and 9 (switched memory B) displayed small negative associations (**Fig. 3C**). Two switched memory B cell clusters (clusters 6 and 10) were increased in current smokers, while the naïve cluster 2 was decreased. Similar associations have been reported before and been linked to chronic inflammation as well as the presence of smoke-induced neo-antigens.^20,25^ Three memory B cell clusters, in particular clusters 6 and 7, displayed significant negative associations with gout, warranting further investigation.

**Fig. 3:**
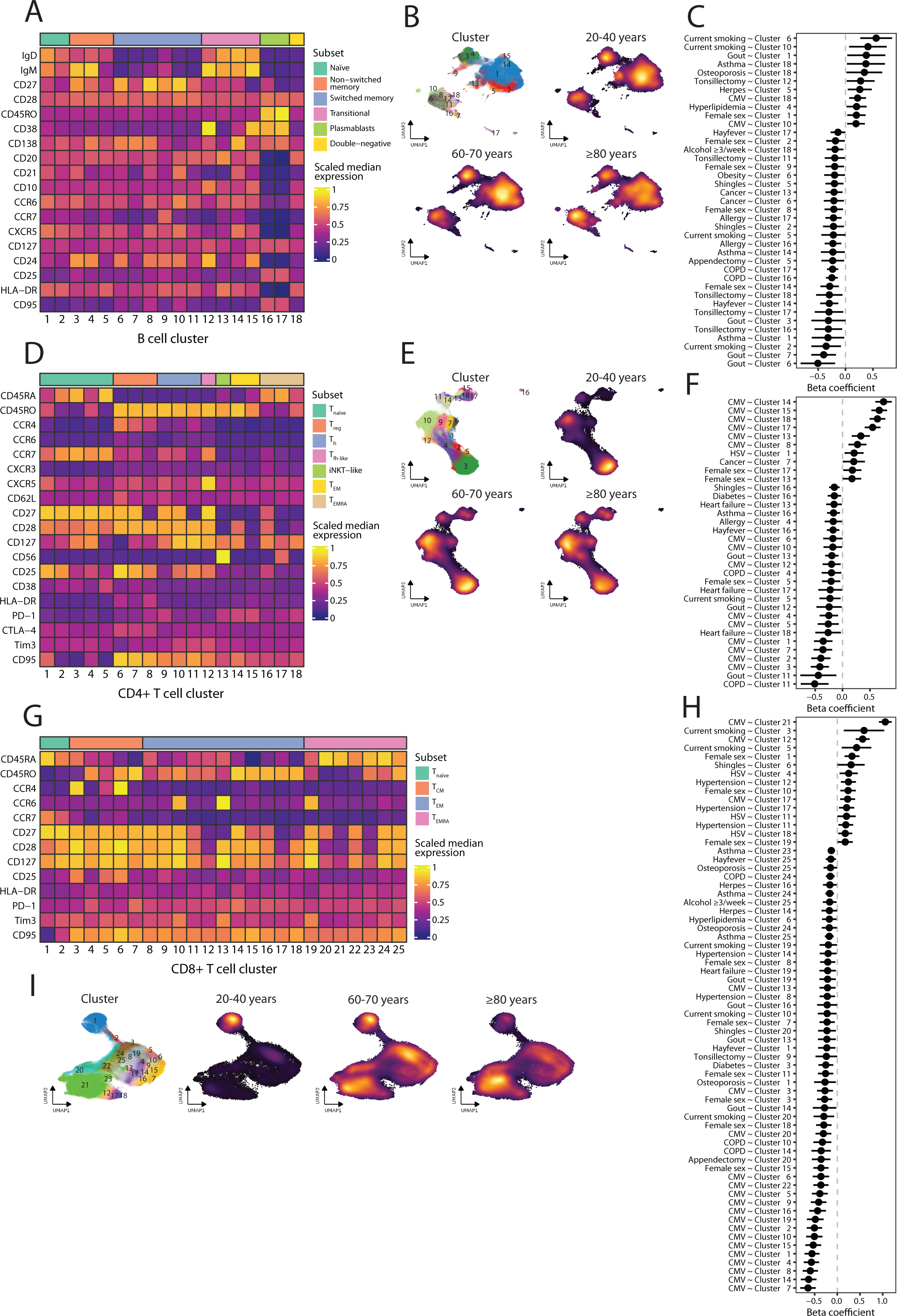
Marker characteristics, age-related changes, and associations of adaptive immune cells with clinical data. **(A, D, G)** Heatmap displaying transformed and row-scaled median marker expressions for each cluster of pre-gated B cells (**A**), CD4+ T cells (**D**), and CD8+ T cells (**G**). Clusters were grouped into sub-lineage subset types based on canonical marker expressions (see methods). (**B**, **E**, **I**) UMAP cluster and density plots depicting the clusters and age distributions of B cells (**B**), CD4+ T cells (**E**), and CD8+ T cells (**H**). (**C, F, H)** Effect size plots with 0.95 confidence intervals from robust linear regression models with scaled cluster frequencies of B cells (**C**), CD4+ T cells (**F**), and CD8+ T cells (**H**) as response, and indicated variables as predictors. All models were adjusted for age and sex, except for models featuring sex, which were only adjusted for age. Only significant predictors (p ≤0.05) are displayed. Abbr.: Allergic disease: presence of at least one allergic disease, CMV: CMV seropositivity, Herpes: history of herpes symptoms (genital or labial), HSV: HSV (type 1 or 2) seropositivity, Shingles: history of shingles, Smoking: history of smoking, Thyroid disease: history of any thyroid disease.

Next, we examined the CD4+ T cell compartment and identified 5 naïve clusters (clusters 1–5), 3 T_reg_ clusters (clusters 6–8), 3 central memory (T_CM_) clusters (clusters 9–11), 1 follicular-helper-like (T_fh-like_) cluster (cluster 12), 1 invariant natural killer T cell (iNKT)-like cluster (cluster 13), 2 T_EM_ cluster (clusters 14–15), and 3 T_EMRA_ clusters (cluster 16–18; **Fig. 3D** and **Extended Data Fig. 6B**). The CD4 T cell compartment displayed substantial changes across age (**Fig. 3E**, **Extended Data Fig. 8**). Only two of the five naïve clusters decreased with age, most prominently the largest cluster 3. All T_reg_ clusters showed frequency increases, as shown previously^14^, while the T_fh-like_ cluster notably decreased within the elderly (**Extended Data Fig. 8, Suppl. Table 6)**. One CD28+ T_EM_ cluster (cluster 14) increased with age, as reported before^26^, while the other T_EM_ and T_EMRA_ clusters remained relatively stable. The iNKT cluster and one T_EMRA_ cluster (cluster 17) displayed subtle positive associations with female sex, while the small naïve cluster 5 tended to have higher frequencies in males (**Fig. 3F**). The strongest associations to clinical or serological data were found for CMV seropositivity. While the differentiated T_EM_ and T_EMRA_ clusters 14, 16, 17, and 18 (but not 16) showed strong positive associations, as did clusters 8 (T_reg_) and 13 (iNKT) to a lesser extent, all naïve clusters and T_reg_ cluster 7 displayed negative associations, although to varying degrees. T_CM_ cluster 11, being CCR4+CCR6-CXCR3-, was negatively associated with COPD and gout.

Lastly, we investigated CD8+ T cells with 25 clusters encompassing 2 naïve clusters (clusters 1–2), 5 T_CM_ clusters (clusters 3–8), 11 T_EM_ clusters (clusters 8–18), and 7 T_EMRA_ (clusters 19–25). Characteristics are displayed in **Fig. 3G** and **Extended Data Fig. 6C**. Similar to the corresponding plot in CD4+ T cells, the UMAP density plot demonstrated strong changes across age groups (**Fig. 3I**). The largest cluster 1 (naïve), displaying a noteworthy high variability in young individuals, was substantially reduced in the elderly, while the frequencies of naïve cluster 2 decreased only marginally with age (**Extended Data Fig. 9**). A particular CCR4+ T_CM_ cluster (cluster 6) substantially increased with age, aligning well with recent single-cell sequencing data describing a similar population that accumulated with age.^14^ Interestingly, those cells increased faster with age compared to CCR4-T_CM_ cells but displayed a higher T cell repertoire clonality.^14^ That study also reported a population of CCR4+HLA-DR^pos^CD27+CD25+ T_EM_ cells, which notably decreased with age, which we also observed in the corresponding cluster 10. Other T_EM_ clusters, such as clusters 7 and 8, clearly increased in elderly individuals. The strongest increase of all CD8+ T cell clusters was observed in T_EMRA_ cluster 21 (**Extended Data Fig. 9, Suppl. Table 7)**. Noteworthy was the high inter-individual frequency variability in this T_EMRA_ cluster, ranging from <5% to >75% of all CD8+ T cells. This cluster also showed the strongest association with CMV seropositivity, which could explain a substantial part of this variability (**Fig. 3H**). Overall, CMV seropositivity displayed a high number of significant associations with cluster frequencies in both directions, affecting clusters from all CD8+ T cell subset types. Moreover, three clusters (clusters 1, 10, and 19) were positively associated with female sex, while clusters 3, 7, 8, 11, 15, and 18 displayed negative associations. We also observed two CCR4+ T_CM_ clusters (clusters 3 and 5) that were positively associated with smoking. Similar populations have been described to be elevated in current smokers by others.^20^

In summary, the high-resolution profiling revealed that the adaptive immune cell composition undergoes pronounced changes with aging. Many age-related changes as well as known and novel associations with clinical and serological data could be pinpointed in more detail to specific subpopulations.

## Multi-block integrative

## analysis of aging

After the detailed study of the innate and adaptive immune cell populations, our aim was to attain a more global picture of the complexity of the aging immune system by integrating different datasets. Specifically, we aimed to decipher distinctive features of aging within the elderly population, rather than solely in contrast to young adults. To identify aging signatures across multiple datasets, we employed a multiple kernel learning based consensus orthogonal partial least square (consensus OPLS) regression analysis to integrate and jointly analyze clinical, laboratory, spectral flow cytometry, and multiplex cytokine data from all elderly participants. Intriguingly, this approach separates systemic variation within the data into predictive components summarizing information related to the outcome variable – in this case, age – and orthogonal components capturing variation uncorrelated to the outcome variable.

The derived model, without including age as an input variable, prominently displayed a discernable age-based ordering of individuals within the score plot along the latent predictive component (**Fig. 4A**), affirming the model’s good predictive ability regarding age using other parameters. Interestingly, the laboratory, spectral flow cytometry, and cytokine datasets contributed nearly equally to the predictive component (31%, 29%, 26%, respectively; **Fig. 4B**). This underscores the rationale to integrate information from different sources, each capturing distinct biological facets, to accurately depict age-related changes in the elderly population. In contrast to the predictive component, the orthogonal components were most strongly influenced by the cytokine data, indicating its significant contribution to variation unrelated to age (**Fig. 4B and C**). In order to identify the key variables driving the predictive performance of the model, we calculated variable importance on projection (VIP) scores, which assess the relative importance of each variable in explaining variation in age in the data. High VIP on the positive side of the predictive component represent features associated with more advanced age in our cohort, while those on the negative predictive component side signify variables characteristic for younger individuals within the elderly cohort. Advanced aging was primarily associated with a set of chemokines including CCL3, CCL27, CXCL 9, HGF, as well as particular immune cell subsets such as B cell cluster 11 (switched memory) and HLA-DR^pos^ cluster 11 (cDC1; **Fig. 4D**). Variables associated with younger elderly individuals were laboratory variables such as platelet and lymphocyte counts, but also several immune cell clusters including CD8 cluster 1 (naïve) and 13 (T_EM_), HLA-DR^pos^ cluster 2 (classical monocyte), and, most prominently, CD4 cluster 12 (T_fh-like_). VIP scores for all relevant variables (i.e., VIP > 1) are provided in **Suppl. Data 1**.

**Fig. 4:**
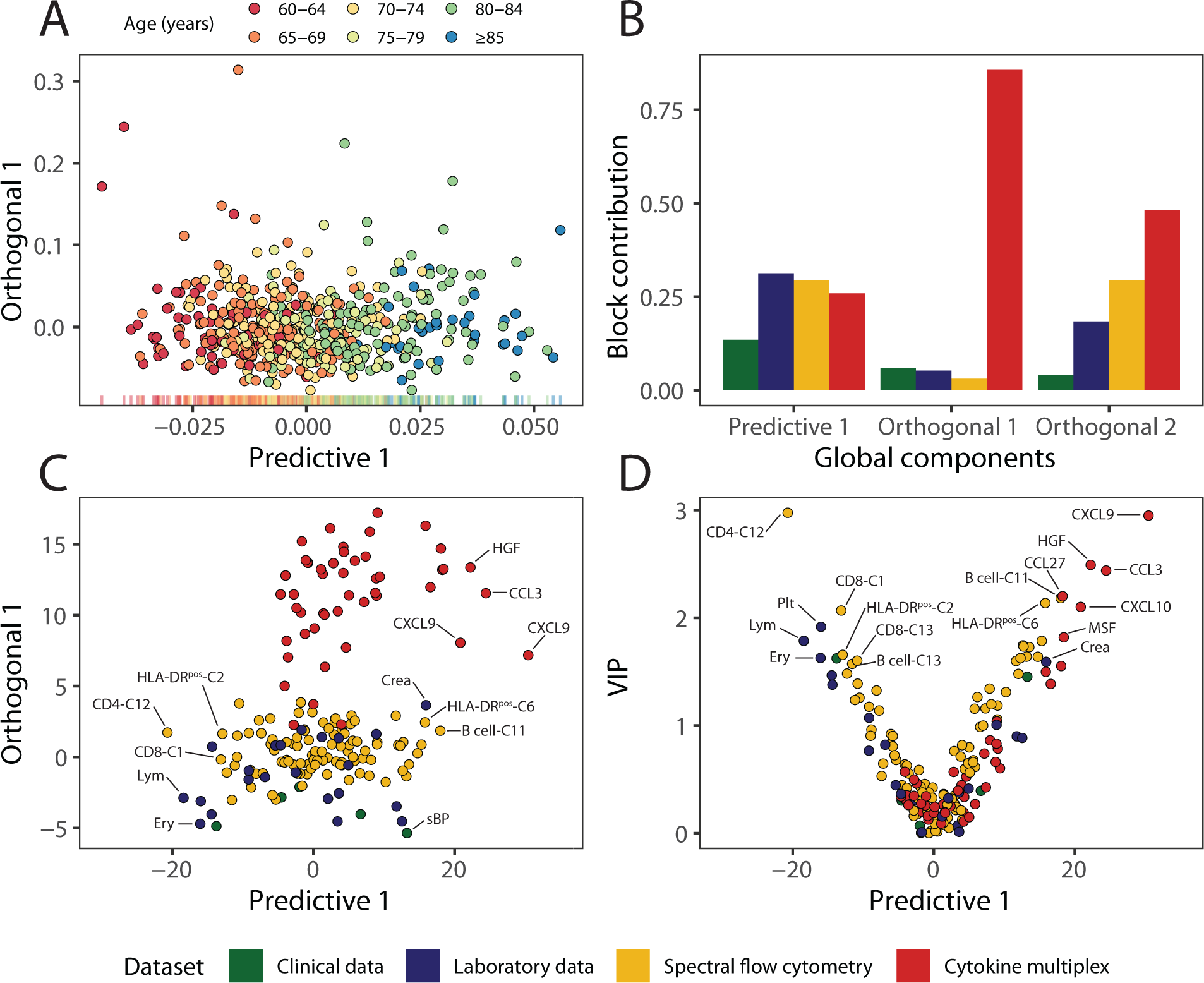
Multi-block integrative analysis of aging. (**A**) Score plot depicting study participants in the new consensus space between (age-related) predictive and (age-unrelated) orthogonal latent components calculated by the integrative consensus OPLS model. (**B)** Contribution plots displaying the weight of each dataset on the predictive and orthogonal components. (**C)** Loading plot showing the projection of variables on the predictive and first orthogonal components. (**D)** Variable Importance on Projection (VIP) plot illustrating VIP coefficients of variables along the predictive component. Higher VIP coefficients indicate a higher relevance of the variable to the predictive ability of the model. Abbr.: ALT: Alanine transaminase, Chol: Cholesterin, Crea: Creatinine, Ery: Erythrocyte counts, Hb: Hemoglobin, HDL: High-density lipoprotein, HR: Heart rate, LDL: Low-density lipoprotein, Leu: Leukocyte counts, Lym: Lymphocyte counts, Mono: Monocyte counts, sBP: Systolic blood pressure, Plt: Platelet counts,

Collectively, these data demonstrate a multifaceted signature of aging, with significant age-related changes occurring across various datasets capturing different aspects of biology. Furthermore, the joint analysis enables to assess the relative importance of different datasets and variables to explain variation in age, which might contribute to a more comprehensive understanding of aging.

## Systems biology perspective on smoking, obesity, and age-related diseases

After finding characteristic features of aging across datasets, we sought to extend this approach to identify integrative signatures of smoking, obesity, and common age-related diseases, including osteoporosis, gout, history of shingles (herpes zoster), and heart failure. To this end, we conducted a consensus OPLS discriminant analysis to delineate factors distinguishing between diseased and non-diseased individuals. In general, distinct group separation was observed for all diseases, albeit often with notable overlap, which is expected given the heterogeneity within diseases and myriad of influencing factors (**Extended Data Fig. 10A-F,** left column). Notably, the contribution of datasets to the predictive component varied significantly across diseases, emphasizing the potential for systems-level perspectives to provide new and complementary insights **(Extended Data Fig. 10A-F,** middle and right column). Detailed predictive component and VIP values for all variables in all analyses are provided in **Suppl. Data 1**.

Laboratory blood counts of leukocytes and their subsets (lymphocytes, monocytes, neutrophils) had the strongest impact on the predictive component for smoking (**Fig. 5A**), consistent with the well-documented phenomenon of smoking-induced leukocytosis.^27,28^ Additionally, several immune cell clusters exhibited high VIP scores. Counterintuitively, CXCL10 was associated with non-smoking, likely due to younger age of smokers in our cohort. In fact, only one out of 55 elderly smokers was older than 81 years, raising speculation that this finding may not be coincidental, given the link between smoking and morbidity^29^, frailty^30^, and physical decline^31^, possibly hindering study visit attendance. Similarly, obesity was most strongly associated with altered laboratory values, with increased levels of uric acid, HbA1c, liver enzymes, and leukocytes, as well as decreased HDL levels (**Fig. 5B**), collectively indicative of an altered metabolic state. We also observed elevated levels of IL-1 receptor antagonist (IL-1RA), which has been linked to increased leptin resistance in obese individuals^32^, while other cytokines and immune clusters displayed weaker associations.

**Figure 5:**
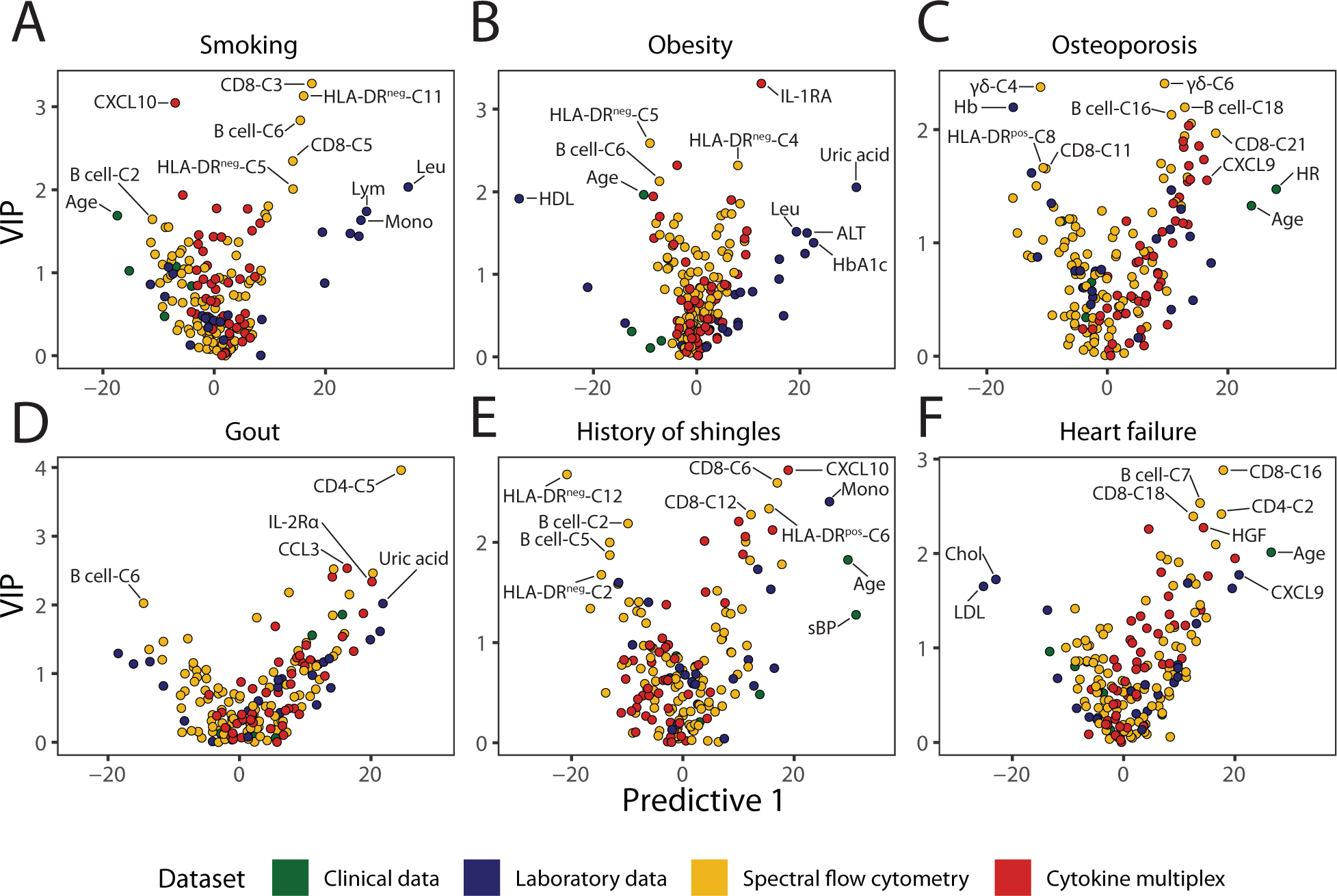
Multi-block integrative analysis of aging and of age-related diseases. VIP plots VIP plots portraying distinguishing features and their relevancy for smoking (**A**), obesity (**B**), as well as osteoporosis (**C**), gout (**D**), participants with a history of shingles (**E**), and heart failure (**F**). Abbr.: ALT: Alanine transaminase, Chol: Cholesterin, Crea: Creatinine, Ery: Erythrocyte counts, Hb: Hemoglobin, HDL: High-density lipoprotein, HR: Heart rate, LDL: Low-density lipoprotein, Leu: Leukocyte counts, Lym: Lymphocyte counts, Mono: Monocyte counts, sBP: Systolic blood pressure, Plt: Platelet counts,

Osteoporosis was strongly associated with age, as expected, and somewhat unexpectedly, increased heart rate (**Fig. 5C**). However, two large studies^33,34^ found heart rate to be an independent predictor of osteoporosis and osteoporotic fractures, in line with our results. Additionally, the integrative analysis revealed associations with several chemokines, such as CXCL9, and immune cell clusters, such as B cell clusters 16 and 18. In gout, uric acid displayed naturally one of the highest predictive component values, while a set of cytokines and immune cell clusters also had a significant predictive impact (**Fig. 5D**). We observed increased levels of soluble IL-2Rα (sCD25) in osteoporosis patients, consistent with a systemic inflammatory state, as well as several naïve CD4 T cell clusters (clusters 1, 2, and 5), which have been found to be enriched in patients with gout before.^35^ In participants with a history of shingles, age displayed expectedly a high predictive value (**Fig. 5E**). Systolic blood pressure and laboratory monocyte counts were also associated with a history of shingles, possibly reflecting the higher age of those participants. Several immune cell clusters, however, also had high predictive component values. CD8 clusters 6 and 12 as well as HLA-DR^pos^ C6 exhibited high positive predictive component values, while HLA-DR^neg^ C2 and 12 displayed negative values. In heart failure, we observed a negative association with cholesterol and LDL levels (**Fig. 5F**). Although these are major predictors for coronary artery disease, patients with advanced heart failure often have low cholesterol levels, possibly due to inflammation and adrenergic activation.^36^ Even the cytokine and immune cell datasets contained variables contributing significantly to the predictive component. Though HGF and CXCL9 might reflect the higher age of patients with heart failure, increased HGF levels have also been specifically linked to heart failure before.^37,38^ Several T cell clusters including CD8 cluster 16 and 18 (T_EM_) and CD4 clusters 2 (naïve) also displayed high predictive component values.

Altogether, the multi-dataset profiling could elucidate systems-level changes in diseased participants by integrating various layers of information. While obesity and smoking were dominated by laboratory variables, the studied diseases often displayed predictive associations with a combination of variables from different datasets. The integrative analysis could potentially lead to important complementary insights when studying those diseases.

## Discussion

Age profoundly influences the human immune landscape.^11^ On the cellular and molecular level, important mechanisms of immunosenescence have been elucidated.^39^ On the systems level, several studies^11,12,14,40^ have delineated significant alterations in the immune cell composition across life, but the understanding of age-related changes in elderly individuals remains incomplete. Studies particularly focusing on elderly individuals are frequently limited in size, leading to inconsistent findings. Moreover, they often solely contrast elderly with young people, overlooking potentially pertinent intra-elderly changes.

Here we performed a comprehensive deep immune profiling study in a well-defined cohort of 550 elderly individuals and 100 young adults, randomly selected from the general population. We used an unsupervised clustering approach to group cells into distinct clusters, a method particularly suitable for analyzing high-dimensional cytometry data. By simultaneously evaluating the entire expression profile in acquired cells, this approach enables the identification of cell clusters in a data-driven and unbiased manner, obviating the need for a manual, iterative gating strategy. As a result, we could often identify multiple distinct clusters within sub-lineage subsets (e.g., within naïve cells). Overall, the unbiased clustering yielding 97 populations allowed us to depict a comprehensive and much more granular picture of the elderly immune cell composition than before. In combination with the large sample size, even nuanced changes within the elderly population could be investigated and were identified for a vast number of subpopulations.

In line with other large-scale studies^11,12,14^, we found high inter-individual variability in numerous cell types, frequently exceeding that observed in smaller studies of elderly individuals. This emphasizes the need for large studies to fully capture the human variability and it is also relevant, for example, when interpreting findings from single-cell sequencing studies, which often only include small numbers of samples due to cost constraints. To a substantial part, the inter-individual variation has been attributed to non-genetic factors.^9,41^ Consistent with other studies from younger cohorts, we found striking associations of several immune cell clusters with known factors such as BMI, sex, and smoking.^11,12,20^ These could be further localized to specific subpopulations; for instance, CMV infection was significantly associated with various clusters of γδ, CD4+ and CD8+ T cells, but the strength of associations varied among different cell clusters within sub-lineage populations. Additionally, the detailed clinical characterization of participants enabled us to test for clinical associations. We identified several clusters with potential associations to specific diseases, such as decreased memory B cell clusters in gout. However, clinical associations often displayed small effect sizes and thus require further validation in additional studies. Contributing to this could be the broad scope of the clinical questionnaire, which did not provide further differentiation regarding disease severity or treatment, potentially resulting in heterogeneous patient groups.

To obtain a global picture of aging and of several common age-related diseases, we used a multi-block integrative analysis approach to merge biological information derived from various datasets. Interestingly, the predictive components for aging and for several age-related diseases were driven by different datasets, with varying weightings. This suggests that an integrated analysis approach can unveil important additional insights. The predictive component for gout, for example, was influenced not only by uric acid, but also by several immune cell clusters and cytokines. A systems biology perspective, which combines different disease aspects, thus may lead to a more holistic understanding, a promising investigative approach for the future. Finally, this approach is particularly adept at generating new research hypotheses, because the joint analysis of a large number of variables can reveal novel or unexpected associations, that may not have been actively sought, such as those observed between osteoporosis and heart rate. These may then stimulate further literature or experimental research.

We note some limitations to our study. Longitudinal variation was not addressed, but has been reported to be low.^11^ Moreover, a longitudinal sub-study of 250 RESIST SI cohort participants is currently underway. Participants were predominately Caucasian and similar studies in other populations are warranted. Other immunological investigations, such as genetics or transcriptomics, were not part of this study, but research efforts addressing them are currently ongoing within the RESIST SI consortium and will complement insights on the elderly immune system.

## Methods

### The RESIST Senior Individuals Cohort

The RESIST SI cohort includes 550 people aged ≥60 years and 100 young adults aged 20–40 years. Study participants were invited by mail to participate in the study after random selection by the local residents’ registry office of Hannover, Germany. Enrollment to the study occurred between December 2019 and March 2022. Inclusion criteria included a permanent residence in Hannover and an age of ≥60 years for the elderly study arm and of 20–40 years for the young adult study arm. Exclusion criteria included the inability to comprehend or provide informed consent, a history of organ transplantation, pregnancy, and immunosuppressive medications (chemotherapy, systemic corticosteroids, cyclophosphamide, methotrexate, azathioprine, biologicals, calcineurin-, mTOR-, JAK-inhibitors). The study was approved by the local ethics committee at Hannover Medical School (8615_BO_S_2019) and written consent was obtained from all participants prior to enrollment. Further details on the RESIST SI cohort have been described elsewhere.^17^

### Clinical data and biomaterials

Detailed information on socio-demographics, behavioral habits, and medical history were obtained by questionnaire at the study visit. The standardized interview was designed to align with that of the German National Cohort^42^, the largest epidemiological study in Germany. Medical history included comorbidities, operations, infection susceptibility, and other pertinent clinical variables. Physical measurements were taken and included, among others, blood pressure, heart rate, weight, waist circumference, and body mass index. Blood samples were obtained and analyzed for blood counts and standard clinical chemistry parameters at the central laboratory at Hannover Medical School. Serological testing was conducted at the Institute of Virology at Hannover Medical School. Various other biomaterials were sampled at the study visit and are described in detail elsewhere.^17^

### Multiplex cytokine assay

Multiplex assays (Bio-Plex Pro Human Cytokine Screening Panel 48-Plex, #12007283 Biorad, USA) were performed on plasma samples of the participants, following the manufacturer’s instructions. Briefly, magnetic capture beads were pipetted into 96-well plates in equal amounts and then washed twice in wash buffer solution. Fifty μL of diluted samples (1:2) and standards were added to the plates and incubated for 30 min at room temperature. After incubation, the plates were washed 3 times in wash buffer solution and detection antibodies were pipetted into each well in equal amounts to be incubated for 30 min at room temperature. After incubation, the plates were once again washed 3 times and a streptavidin-phycoerythrin solution was added to each well in equal amounts for 10 minutes incubation at room temperature. Plates were then washed thrice for the last time and stored at 4° before measurement in a Bio-Plex 200 System (#171000201 Biorad, USA). All samples were prepared with the same reagents on the same day and were measured together the next day. Four samples were excluded from the analysis due to technical sampling errors arising from the presence of erythrocytes in the plasma samples. For analysis, measured concentrations below the lower limit of detection were set to half the minimum value of the standard curve of the respective analyte. In total, 642 (99%) samples were available for measurement and down-stream analyses.

### Spectral flow cytometry

Whole blood samples were collected from study participants into CPT tubes (BD Biosciences) and processed into peripheral blood mononuclear cells (PBMCs), as described by the manufacturer. As expected, the resulting samples contained no neutrophilic granulocytes after PBMC isolation, as confirmed by flow cytometry. Samples were stained with two different antibody panels (**Supplementary Table 8-9**) to investigate innate and adaptive immune cell subsets, respectively, as well as the surface expression of various activation and exhaustion markers. After thawing, samples were washed twice with PBS, stained at room temperature for 20 min and then washed twice again with PBS before acquisition. Samples were then acquired on a spectral flow cytometer (Cytek Aurora) equipped with five lasers operating at 355 nm, 405 nm, 488 nm, 561 nm, and 640 nm. Data were acquired using SpectroFlow (Cytek, version 3.0.3). Dead cells and cell doublets were removed from the data using FCS Express 7 (Denovo Software). For the high-resolution clustering analysis, cells were pre-gated on HLA-DR^neg^ innate cells (CD45+CD3-CD19-HLA-DR^neg^), HLA-DR^pos^ innate cells (CD45+CD3-CD19-HLA-DR^pos^), γδ T cells (CD45+CD3+TCRγδ+), B cells (CD45+CD19+), CD4 T cells (CD45+CD3+CD4+), and CD8 T cells (CD45+CD3+CD8+) in FCS Express prior to the clustering analysis (**Extended Data** Fig. 1). The pre-gated files of each cell lineage were exported as FCS files from FCS Express for further analysis. In total, 646 (99%) and 644 (99%) samples were available for analysis for the innate and adaptive immune cell panel, respectively.

### High-dimensional flow cytometry data analysis

FCS files were loaded in R and transformed using the asinh transformation in the flowVS package^43^. Batch-correction accounting for the different time points of acquisition was performed using the CyCombine algorithm^44^ with the normalization method set to “rank” and a 10×10 grid size. After batch-correction, the data from each pre-gated lineage were visually inspected using multidimensional scaling plots and marker expression plots stratified by batch to confirm the absence of batch-effects before proceeding with the analysis. Subsequently, we employed the unsupervised Flow Self-Organizing Maps (FlowSOM) algorithm^45^ to assign cells to cell clusters based on their global marker expressions. Within each pre-gated lineage, surface markers known to distinctively separate cell subtypes were chosen for clustering, based on previous literature^46–48^ and experiences in our group^49–52^. Activation markers were not used for clustering as activated cells clustered together independent of their subtype, but were displayed in heatmaps^53^ and ridgeplots and could consequently be used for cluster annotation. For HLA-DR^neg^ innate cells and HLA-DR^pos^ innate cells, the following markers were used, respectively: CD16, CD11c, CD56, CD86, CD123, CD103, CCR5, HLA-DR, CD80, CD127, NKG2D, CRTH2, CCR2, CD66b, CD64, CD206 (HLA-DR^neg^), and CD16, CD14, CD115, CD11c, CD56, CD86, CD123, CD103, CCR5, HLA-DR, CD141, CD125, CD1c, NKG2D, CRTH2, CCR2, CD66b, and CD64 (HLA-DR^pos^). In HLA-DR^neg^ innate cells, four small clusters of left-over monocytes (HLA-DR^pos^CD64+CD16-CD11c+) were removed before the remaining cells were re-clustered. Similarly, in HLA-DR^pos^ innate cells, one small cluster of left-over neurophilic granulocytes (CD66b+) and another small cluster of left-over NK cells (CD56+NKG2D+HLA-DR^low^) were removed and then re-clustered. For B cells, CD21, CD27, IgD, CD138, CD38, and IgM were used. For CD4+ T cells and CD8+ T cells, CD45RA, CCR4, CD27, CD45RO, CCR6, CCR7 were used. γδ T cells were clustered using CD45RA, CD16, CD4, CCR4, CD56, Vg9, CCR6, and CCR7. As described before^45,54,55^, we followed an “over-clustering” approach in order not to miss rare, but distinct cell subpopulations, by firstly setting the number of FlowSOM clusters much higher than the expected cell subtypes. After exploring and comparing the characteristics of those clusters, highly similar clusters were then manually merged, based on biological knowledge of the cell subtypes and markers. The final clusters of each pre-gated lineage were then annotated based on the marker distributions and re-ordered into subset types based on canonical marker expressions. HLA-DR^neg^ innate cells were ordered into early NK cells (CD56^high^CD16^low^), mature NK cells (CD56+CD16+NKG2D+), terminal NK cells (CD16+NKG2D+), CD56-CD16-NKs (CD56-CD16-NKG2D+), basophils (CD123+CRTH2+CCR2+), mast cells (Lin-CD11c^low^CD123^low^), ILCs (Lin-CD16-CD127+) and one undefined cluster. HLA-DR^pos^ innate cells were grouped into classical monocytes (CD14+CD16-), intermediate monocytes (CD14+CD16+), non-classical monocytes (CD14-CD16+), granulo-monocytic precursors (CD16^low^CD11c^low^CCR2+), pDCs (CD123+CCR2+CCR5+), cDC1 (CD11c+CD141+), and cDC2 (CD11c^high^CCR2+CCR5+CD1c+), as well as one undefined cluster. γδ T cells were grouped into Vγ9-Naïve (Vγ9-CD45RA+CD62L^high^CD16-), Vγ9-T_EM_ (Vγ9-CD45RA+CD62LlowCD16^mid^), Vγ9+ Naïve (Vγ9+CD45RA+CD27+CD28+/-CD127+), Vγ9+ T_CM_ (Vγ9+CD45RA-CD45RO+CD27+), Vγ9+ T_EM_ (Vγ9+CD45RA-CD45RO+CD27-), Vγ9+ T_EMRA_ (Vγ9+CD45RA+CD27-CD16+CD56^mid^), and one undefined cluster. B cells were divided into naïve cells (CD27-IgD+), non-switched memory cells (CD27+IgD+), switched memory cells (CD27+IgD-), translational B cells (defined by IgM, CD38, and CD10), plasmablasts (CD20-CD38+CD45RO+), and double-negative cells (IgD-IgM-CD27-). CD4+ T cells were assigned to naïve cells (CD45RA+CCR7+), T_regs_ (CD25+CD127-), T_CM_ (CD45RO+CCR7+ and cytokine receptors CCR4, CCR6, and CXCR3), T_fh-like_ (CD45RO+CCR7+CXCR5+), iNKT (CD45RO+CCR7-CD56+), T_EM_ (CD45RO+CCR7-CD27-), and T_EMRA_ (CD45RA+CCR7-). CD8+ T cells were grouped to naïve cells (CD45RA+CCR7+), T_CM_ (CD45RO+ to CD45RO^mid^ and CCR7+ to CCR7^low^), T_EM_ (CD45RA-CCR7-), and T_EMRA_ (CD45RA+CCR7-). Dimensionality reduction was conducted using the Uniform Manifold Approximation and Projection (UMAP) approach. Heatmaps displaying the marker expression profile of clusters within each population were generated^53^ by calculating the row-wise scaled median expression of relevant markers.

### Multi-block integration and analysis

We employed a multi-block consensus orthogonal partial least square analysis (consensus OPLS) for multi-dataset integration and analysis. Consensus OPLS is a low-level data fusion approach allowing to jointly analyze multiple datasets using multiple kernel learning.^56^ The model is limited to continuous input variables, but is robust to different numbers of variables per dataset.^56^ Four datasets were integrated, namely clinical data (age, BMI, waist circumference, heart rate, systolic and diastolic blood pressure), laboratory data (22 standard laboratory values), spectral flow cytometry data (97 cell clusters), and cytokine data (48 cytokines). One laboratory variable, C-reactive protein (CRP), was not available in 111 participants (20%), and therefore excluded from the consensus OPLS-DA modeling, which requires complete observations. All remaining variables were available in >97% of participants. Missing values in those variables were imputed using Random Forest single imputation^57^, yielding a complete dataset of 550 observations. Before modeling, variables were scaled to unit variance and centered to a mean of zero. For the investigation of aging features within the elderly population, we used consensus OPLS regression analysis with age in years as outcome variable. For the analysis of disease-associated features, we used the variant consensus OPLS discriminant analysis with diseased vs. non-diseased as outcome groups. For the obesity model, BMI and waist circumference were removed as input variables, as they define obesity. For all models, the optimal number of latent predictive and orthogonal variables was determined based on the DQ^2^ index, estimated using 10-fold Cross Validation. The importance of each variable in explaining variability regarding age and disease separation was assessed using the Variable Importance on Projection (VIP) metric.^58^

### Statistics

Demographical, clinical, and serological data were summarized using descriptive statistics. Group comparisons between two groups were conducted using the Wilcoxon test. Two-sided non-parametric Jonckheere-Terpstra tests were used to test for trends in cluster frequencies across age groups. To test for associations between cluster frequencies and clinical variables, linear regression models with robust standard errors^59^ were constructed with adjustment for age and sex, except for models testing sex, which were only adjusted for age. Clinical variables were dichotomized as independent variables, and cluster frequencies were scaled to unit variance and centered as dependent variables. Correlations between cytokine levels and cluster frequencies were evaluated using Spearman’s rank correlation. The statistical analysis used in each figure is specified in the respective figure legend. P values ≤0.05 were considered statistically significant and were not adjusted because of the exploratory, non-confirmatory nature of this study.^60^ All statistical analyses were conducted with the statistical software R (version 4.3.2).^61^

### Data availability

Source data is provided with this paper. Raw data can be accessed by researchers who provide a written study proposal, which has to be approved by the RESIST SI Cohort steering committee.

### Code availability

Computer code used in this study will be made freely available upon final acceptance of the paper.

## Acknowledgements

This study was funded by the Deutsche Forschungsgemeinschaft (DFG, German Research Foundation) under Germany’s Excellence Strategy – EXC 2155 – project number 390874280. LR was supported by the Hannover Biomedical Research School (HBRS), the Center for Infection Biology (ZIB), the Else Kröner-Fresenius Stiftung (2018_Kolleg.12), and the Joachim Herz Stiftung.

## Author information

### Consortium

The RESIST SI Cohort Investigators consist of the following persons in addition to the authors of this study: Berislav Bošnjak, Institute of Immunology, Hannover Medical School (MHH), Hannover, Germany; Felix Jenniches, Department for Epidemiology, Helmholtz Centre for Infection Research (HZI), Braunschweig, Germany; Norman Klopp, Hannover Unified Biobank (HUB), Hannover Medical School (MHH), Hannover, Germany; Till Robin Lesker, Department of Microbial Immune Regulation, Helmholtz Centre for Infection Research (HZI), Braunschweig, Germany; Martin Stangel, Department of Neurology, Hannover Medical School, Hannover (MHH), Germany, and Department of Translational Medicine Neuroscience, Novartis Institute for BioMedical Research, Basel, Switzerland.

### Contributions

Study design: T.F.S., G.H., T.W., R.F, L.Ro. Experiments: R.G., I.O., J.B-M. Data analysis: L.Ri, R.G. Data interpretation: L.Ri., R.G., I.O., J.B-M., L.Ro, X. L-L, T.F.S., G.H., T.W, R.F., Cytokine multiplex interpretation: C.F. Writing: L.Ri., R.F. Editing and revising: R.G., I.O., J.B-M., L.Ro, X. L-L, T.F.S., G.H., T.W. Supervision: R.F.

## Competing interest declaration

The authors declare no competing interests.

**Extended Data Fig. 1:** Gating strategy of innate and adaptive immune cells prior to computational clustering (pre-gating).

**Extended Data Fig. 2:** UMAP feature plots of HLA-DR^neg^ innate cells (**A**), HLA-DR^pos^ innate cells (**B**), and γδ T cells (**C**).

**Extended Data Fig. 3:** Scatter plots displaying cluster frequencies of HLA-DR^neg^ innate cells across age groups. The red crossbar indicates the median.

**Extended Data Fig. 4:** Scatter plots displaying cluster frequencies of HLA-DR^pos^ innate cells across age groups. The red crossbar indicates the median.

**Extended Data Fig. 5:** Scatter plots displaying cluster frequencies of γδ T cells across age groups. The red crossbar indicates the median.

**Extended Data Fig. 6:** UMAP feature plots of B cells (**A**), CD4+ T cells (**B**), and CD8+ T cells (**C**).

**Extended Data Fig. 7:** Scatter plots displaying cluster frequencies of B cells across age groups. The red crossbar indicates the median.

**Extended Data Fig. 8:** Scatter plots displaying cluster frequencies of CD4+ T cells across age groups. The red crossbar indicates the median.

**Extended Data Fig. 9:** Scatter plots displaying cluster frequencies of CD8+ T cells across age groups. The red crossbar indicates the median.

**Extended Data Fig. 10:** Score plots (left column), contribution plots (middle column), and loading plots (right column) of consensus OPLS discriminant analysis models for smoking (**A**), obesity (**B**), osteoporosis (**C**), gout (**D**), history of shingles (**E**), heart failure (**F**).

## References

1. United Nations. UN Decade of Heathy Ageing: Plan of Action. https://www.who.int/publications/m/item/decade-of-healthy-ageing-plan-of-action (2020).

2. World Health Organization. WHO Fact Sheet: Ageing and Health. https://www.who.int/news-room/fact-sheets/detail/ageing-and-health (2022).

3. Weyand, C. M. & Goronzy, J. J. Aging of the Immune System. Mechanisms and Therapeutic Targets. Ann Am Thorac Soc 13, S422–S428 (2016).

4. Yoshikawa, T. T. Epidemiology and Unique Aspects of Aging and Infectious Diseases. Clinical Infectious Diseases 30, 931–933 (2000).

5. White, M. C., Holman, D. M., Goodman, R. A. & Richardson, L. C. Cancer Risk Among Older Adults: Time for Cancer Prevention to Go Silver. Gerontologist 59, S1–S6 (2019).

6. Ciabattini, A. et al. Vaccination in the elderly: The challenge of immune changes with aging. Semin Immunol 40, 83–94 (2018).

7. Bonaguro, L. et al. A guide to systems-level immunomics. Nat Immunol 23, 1412–1423 (2022).

8. Brodin, P. & Davis, M. M. Human immune system variation. Nat Rev Immunol 17, 21–29 (2017).

9. Brodin, P. et al. Variation in the Human Immune System Is Largely Driven by Non-Heritable Influences. Cell 160, 37–47 (2015).

10. Roederer, M. et al. The Genetic Architecture of the Human Immune System: A Bioresource for Autoimmunity and Disease Pathogenesis. Cell 161, 387–403 (2015).

11. Carr, E. J. et al. The cellular composition of the human immune system isshaped by age and cohabitation. Nat Immunol 17, 461–468 (2016).

12. Patin, E. et al. Natural variation in the parameters of innate immune cells is preferentially driven by genetic factors. Nat Immunol 19, 302–314 (2018).

13. Mathew, D. et al. Deep immune profiling of COVID-19 patients reveals distinct immunotypes with therapeutic implications. Science (1979) 369, eabc8511 (2020).

14. Terekhova, M. et al. Single-cell atlas of healthy human blood unveils age-related loss of NKG2C+GZMB−CD8+ memory T cells and accumulation of type 2 memory T cells. Immunity 56, 2836–2854.e9 (2023).

15. Zheng, H. et al. Multi-cohort analysis of host immune response identifies conserved protective and detrimental modules associated with severity across viruses. Immunity 54, 753–768.e5 (2021).

16. Saint-André, V. et al. Smoking changes adaptive immunity with persistent effects. Nature 626, 827–835 (2024).

17. Roesner, L. M. et al. The RESIST Senior Individuals Cohort: Design, participant characteristics and aims. medRxiv 2024.04.29.24306533 (2024) doi:10.1101/2024.04.29.24306533.

18. Woelfle, T. et al. Health impact of seven herpesviruses on (pre)diabetes incidence and HbA1c: results from the KORA cohort. Diabetologia 65, 1328–1338 (2022).

19. Franceschi, C., et al. Inflamm-aging: An Evolutionary Perspective on Immunosenescence. Ann N Y Acad Sci 908, 244–254 (2000).

20. Piaggeschi, G. et al. Immune Trait Shifts in Association With Tobacco Smoking: A Study in Healthy Women. Front Immunol 12, (2021).

21. Argentati K et al. Numerical and functional alterations of circulating gammadelta T lymphocytes in aged people and centenarians. J Leukoc Biol 72(**1**), 65–71 (2002).

22. Wistuba-Hamprecht, K., Frasca, D., Blomberg, B., Pawelec, G. & Derhovanessian, E. Age-associated alterations in γδ T-cells are present predominantly in individuals infected with Cytomegalovirus. Immunity & Ageing 10, 26 (2013).

23. Caccamo, N., Dieli, F., Wesch, D., Jomaa, H. & Eberl, M. Sex-specific phenotypical and functional differences in peripheral human Vγ9/Vδ2 T cells. J Leukoc Biol 79, 663–666 (2006).

24. Pitard, V. et al. Long-term expansion of effector/memory Vδ2− γδ T cells is a specific blood signature of CMV infection. Blood 112, 1317–1324 (2008).

25. Brandsma, C.-A. et al. Increased levels of (class switched) memory B cells in peripheral blood of current smokers. Respir Res 10, 108 (2009).

26. Vallejo, A. N. CD28 extinction in human T cells: altered functions and the program of T-cell senescence. Immunol Rev 205, 158–169 (2005).

27. Pedersen, K. M. et al. Smoking and Increased White and Red Blood Cells. Arterioscler Thromb Vasc Biol 39, 965–977 (2019).

28. Jayasuriya, N. A. et al. Smoking, blood cells and myeloproliferative neoplasms: meta-analysis and Mendelian randomization of 2·3 million people. Br J Haematol 189, 323–334 (2020).

29. LaCroix, A. Z. & Omenn, G. S. Older Adults and Smoking. Clin Geriatr Med 8, 69–88 (1992).

30. Kojima, G., Iliffe, S. & Walters, K. Smoking as a predictor of frailty: a systematic review. BMC Geriatr 15, 131 (2015).

31. Rapuri, P. B., Gallagher, J. C. & Smith, L. M. Smoking Is a Risk Factor for Decreased Physical Performance in Elderly Women. The Journals of Gerontology: Series A 62, 93–99 (2007).

32. Meier, C. A. et al. IL-1 Receptor Antagonist Serum Levels Are Increased in Human Obesity: A Possible Link to the Resistance to Leptin? J Clin Endocrinol Metab 87, 1184–1188 (2002).

33. Kado, D. M., Lui, L. L., Cummings, S. R. & Group, T. S. of O. F. R. Rapid Resting Heart Rate: A Simple and Powerful Predictor of Osteoporotic Fractures and Mortality in Older Women. J Am Geriatr Soc 50, 455–460 (2002).

34. Jung, M.-H., Youn, H.-J., Ihm, S.-H., Jung, H. O. & Hong, K.-S. Heart Rate and Bone Mineral Density in Older Women with Hypertension: Results from the Korea National Health and Nutritional Examination Survey. J Am Geriatr Soc 66, 1144–1150 (2018).

35. Chang, J.-G. et al. Single-cell RNA sequencing of immune cells in patients with acute gout. Sci Rep 12, 22130 (2022).

36. Velavan, P., Huan Loh, P., Clark, A. & Cleland, J. G. F. The Cholesterol Paradox in Heart Failure. Congestive Heart Failure 13, 336–341 (2007).

37. Ferraro, R. A. et al. Hepatocyte Growth Factor and Incident Heart Failure Subtypes: The Multi-Ethnic Study of Atherosclerosis (MESA). J Card Fail 27, 981–990 (2021).

38. Lamblin, N. et al. Prognostic significance of circulating levels of angiogenic cytokines in patients with congestive heart failure. Am Heart J 150, 137–143 (2005).

39. Liu, Z. et al. Immunosenescence: molecular mechanisms and diseases. Signal Transduct Target Ther 8, 200 (2023).

40. Jalali, S. et al. A high-dimensional cytometry atlas of peripheral blood over the human life span. Immunol Cell Biol 100, 805–821 (2022).

41. Orrù, V. et al. Genetic Variants Regulating Immune Cell Levels in Health and Disease. Cell 155, 242–256 (2013).

42. Peters, A. et al. Framework and baseline examination of the German National Cohort (NAKO). Eur J Epidemiol 37, 1107–1124 (2022).

43. Azad, A., Rajwa, B. & Pothen, A. flowVS: channel-specific variance stabilization in flow cytometry. BMC Bioinformatics 17, 291 (2016).

44. Pedersen, C. B. et al. cyCombine allows for robust integration of single-cell cytometry datasets within and across technologies. Nat Commun 13, 1698 (2022).

45. Van Gassen, S. et al. FlowSOM: Using self-organizing maps for visualization and interpretation of cytometry data. Cytometry Part A 87, 636–645 (2015).

46. Staser, K. W., Eades, W., Choi, J., Karpova, D. & DiPersio, J. F. OMIP-042: 21-color flow cytometry to comprehensively immunophenotype major lymphocyte and myeloid subsets in human peripheral blood. Cytometry Part A 93, 186–189 (2018).

47. Moncunill, G., Han, H., Dobaño, C., McElrath, M. J. & De Rosa, S. C. OMIP-024: Pan-leukocyte immunophenotypic characterization of PBMC subsets in human samples. Cytometry Part A 85, 995–998 (2014).

48. Baumgart, S., Peddinghaus, A., Schulte-Wrede, U., Mei, H. E. & Grützkau, A. OMIP-034: Comprehensive immune phenotyping of human peripheral leukocytes by mass cytometry for monitoring immunomodulatory therapies. Cytometry Part A 91, 34–38 (2017).

49. Odak, I. et al. Systems biology analysis reveals distinct molecular signatures associated with immune responsiveness to the BNT162b COVID-19 vaccine. EBioMedicine 99, (2024).

50. Odak, I. et al. Brief research report: in-depth immunophenotyping reveals stability of CD19 CAR T-cells over time. Front Immunol 15, (2024).

51. Odak, I. et al. Spectral flow cytometry cluster analysis of therapeutic donor lymphocyte infusions identifies T cell subsets associated with outcome in patients with AML relapse. Front Immunol 13, (2022).

52. Barros-Martins, J., Bruni, E., Fichtner, A. S., Cornberg, M. & Prinz, I. OMIP-084: 28-color full spectrum flow cytometry panel for the comprehensive analysis of human γδ T cells. Cytometry Part A 101, 856–861 (2022).

53. Gu, Z., Eils, R. & Schlesner, M. Complex heatmaps reveal patterns and correlations in multidimensional genomic data. Bioinformatics 32, 2847–2849 (2016).

54. Nowicka, M. et al. CyTOF workflow: differential discovery in high-throughput high-dimensional cytometry datasets. F1000Res (2019).

55. Saeys, Y., Van Gassen, S. & Lambrecht, B. N. Computational flow cytometry: helping to make sense of high-dimensional immunology data. Nat Rev Immunol 16, 449–462 (2016).

56. Boccard, J. & Rutledge, D. N. A consensus orthogonal partial least squares discriminant analysis (OPLS-DA) strategy for multiblock Omics data fusion. Anal Chim Acta 769, 30–39 (2013).

57. Stekhoven, D. J. & Bühlmann, P. MissForest—non-parametric missing value imputation for mixed-type data. Bioinformatics 28, 112–118 (2012).

58. Mahieu, B., Qannari, E. M. & Jaillais, B. Extension and significance testing of Variable Importance in Projection (VIP) indices in Partial Least Squares regression and Principal Components Analysis. Chemometrics and Intelligent Laboratory Systems 242, 104986 (2023).

59. Blair, G., Cooper, J., Coppock, A., Humphreys, M. & Sonnet, L. estimatr: Fast Estimators for Design-Based Inference. 2024 (2024).

60. Rothman, K. J. No Adjustments Are Needed for Multiple Comparisons. Epidemiology 1, 43–46 (1990).

61. R Core Team. R: A Language and Environment for Statistical Computing. Preprint at https://www.r-project.org/ (2024).

