## Supplementary material for "Integrative deep immune profiling of the elderly reveals systems-level signatures of aging, sex, smoking, and clinical traits": Figure S1

Gating Strategy for the innate cell antibody panel:

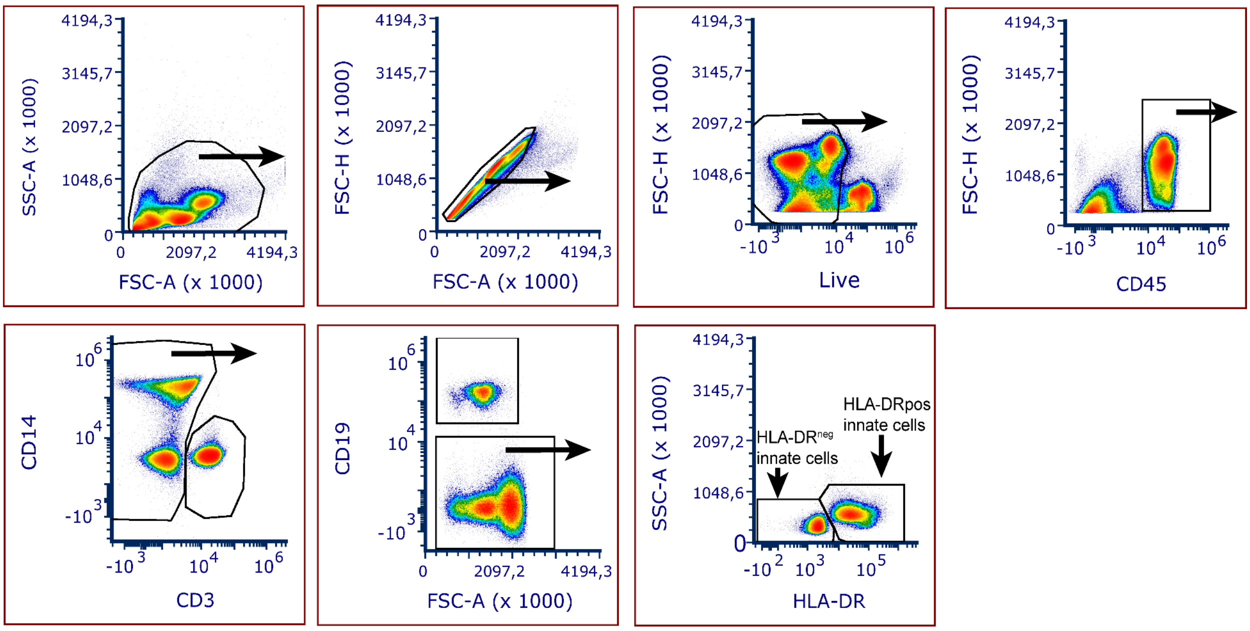


Gating Strategy for the adaptive cell antibody panel:

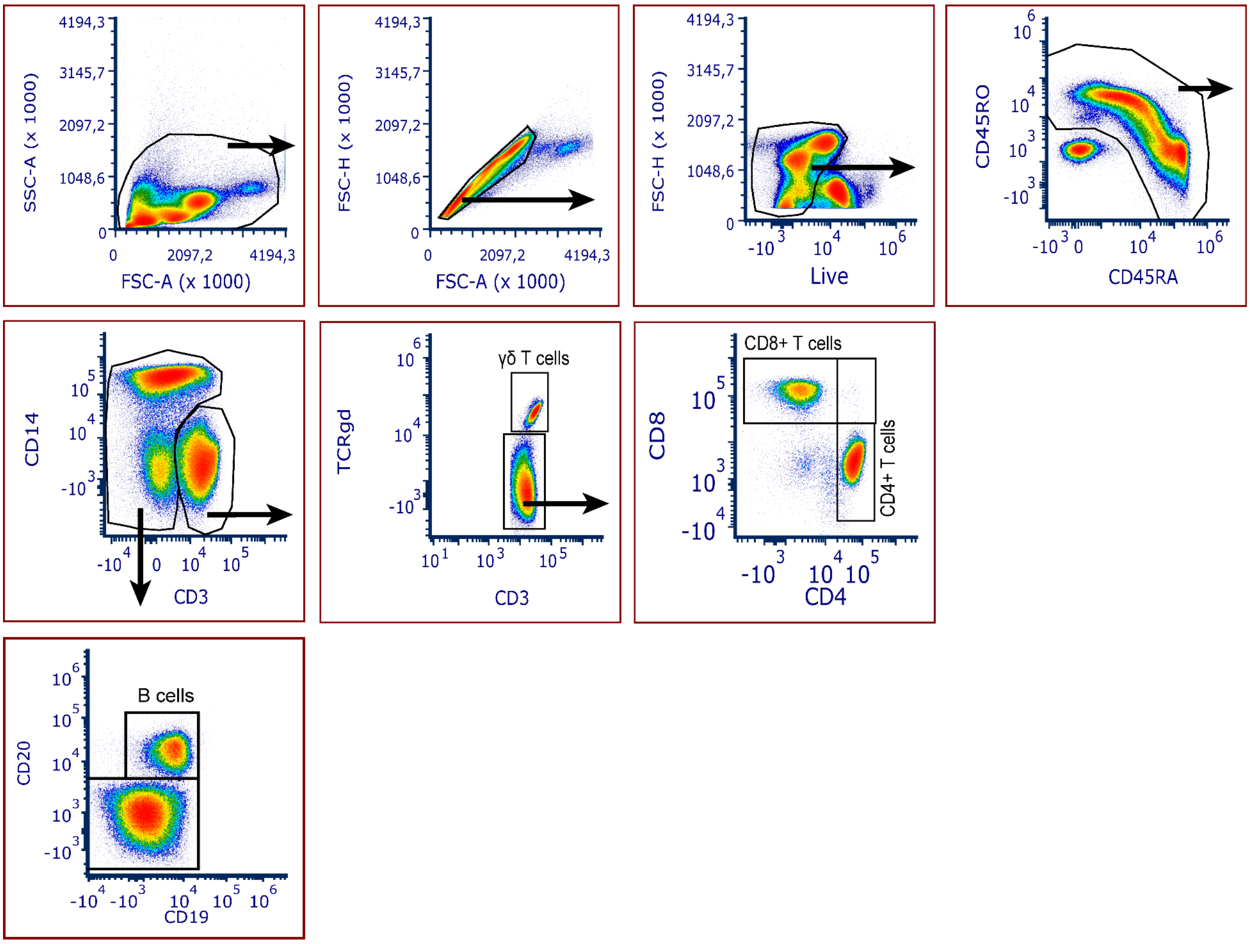
