## Supplementary figures and images for "Integrative deep immune profiling of the elderly reveals systems-level signatures of aging, sex, smoking, and clinical traits"

### Figure S2

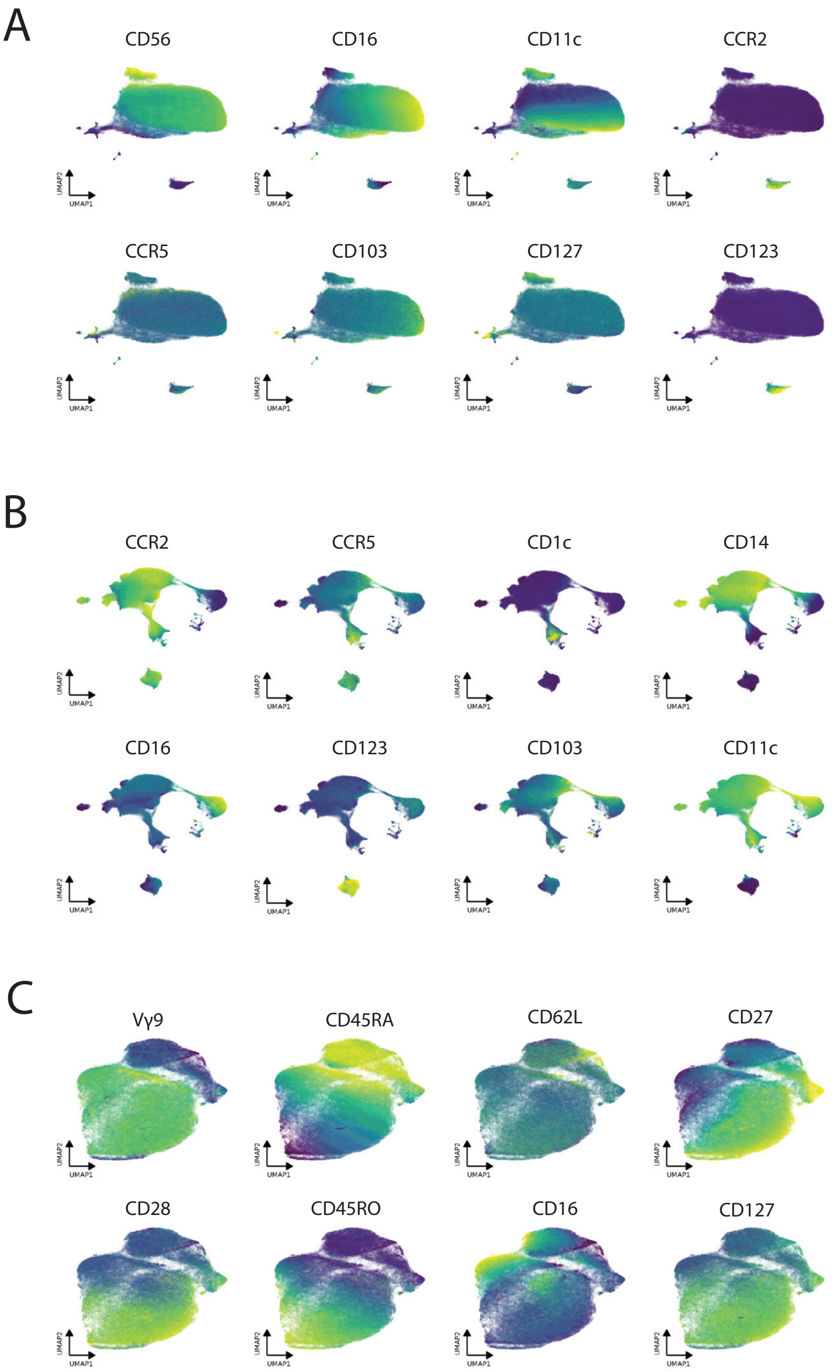

### Figure S3

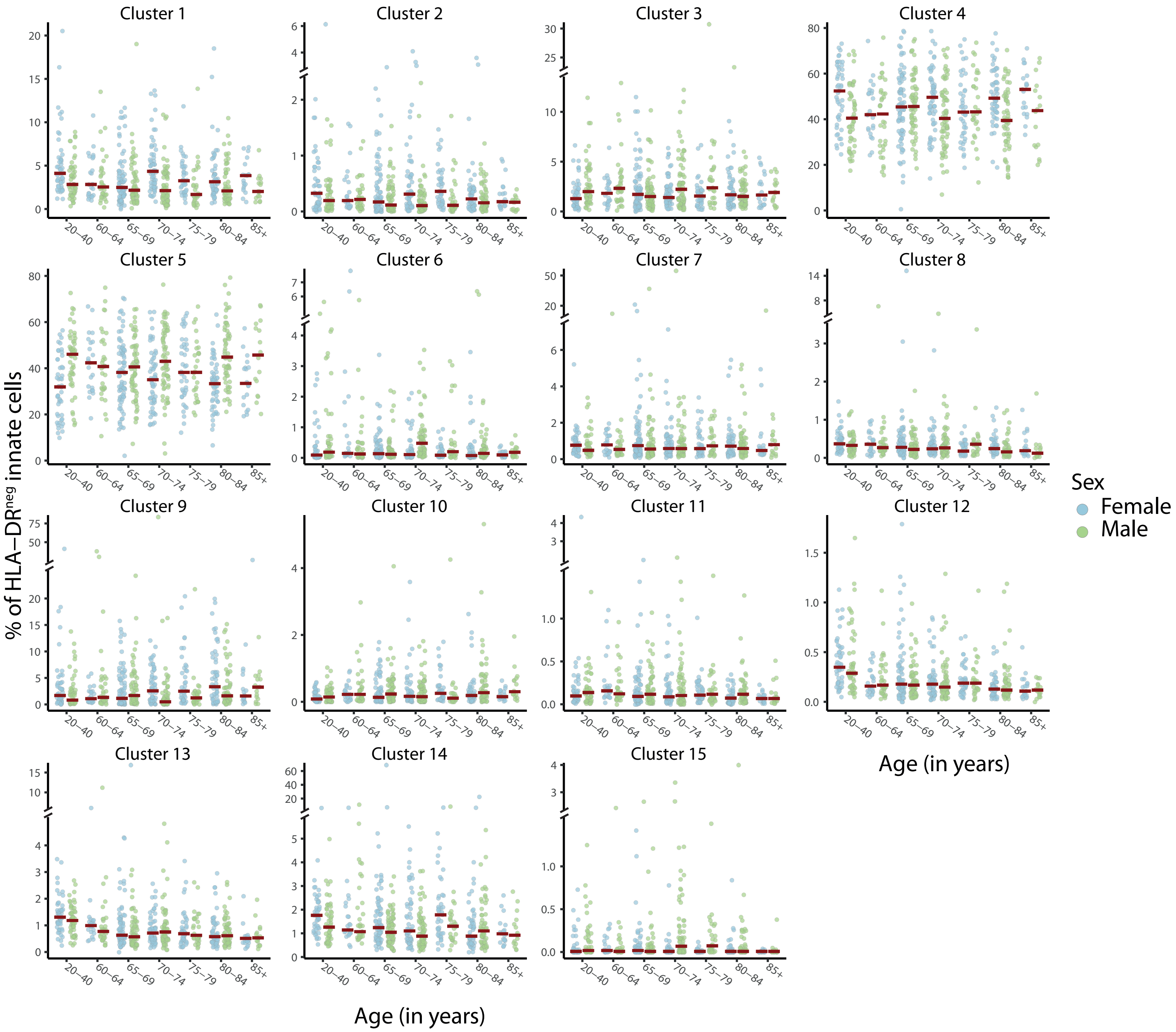

### Figure S4

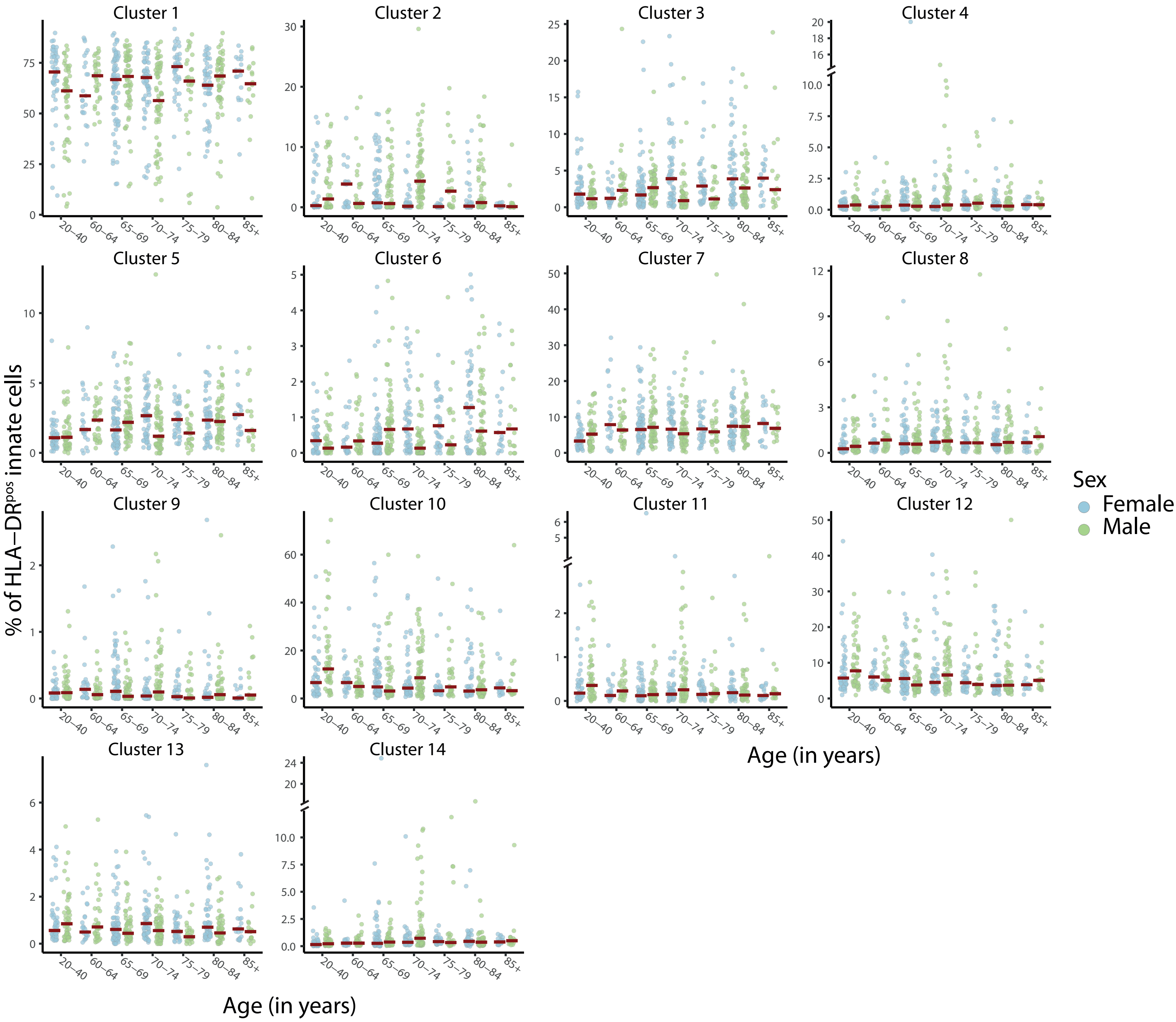

### Figure S5

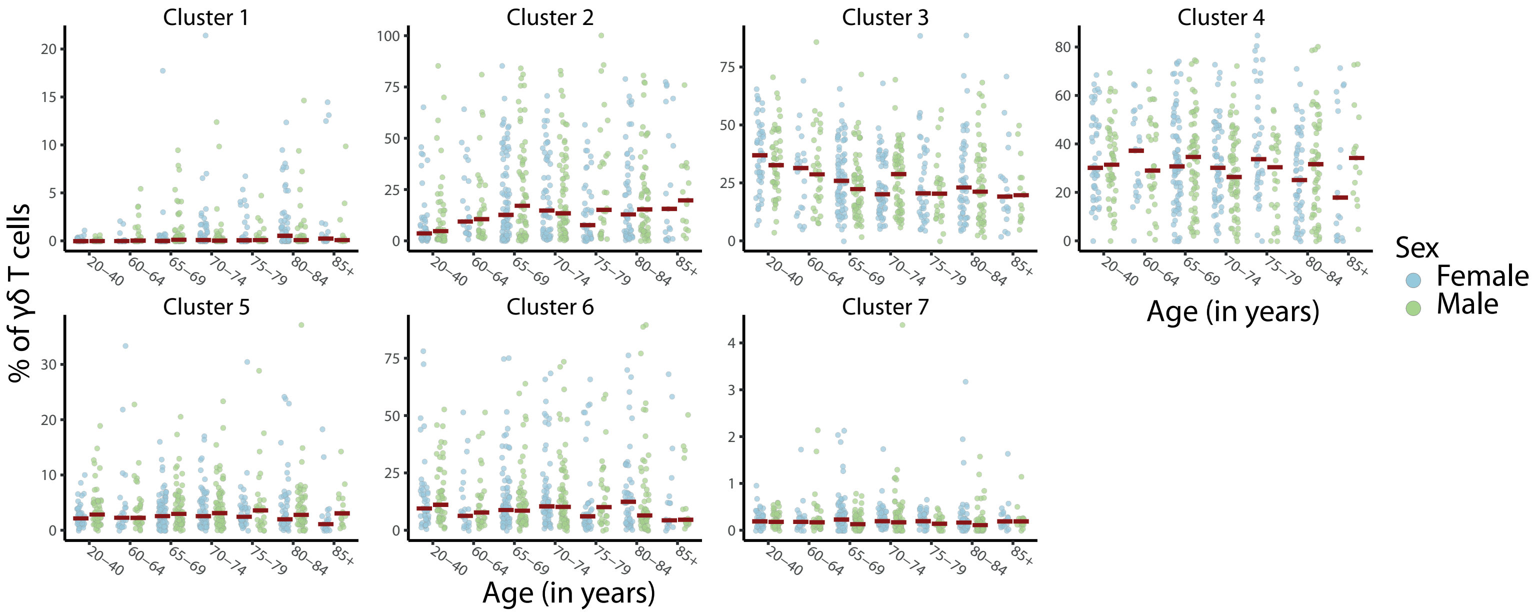

### Figure S6

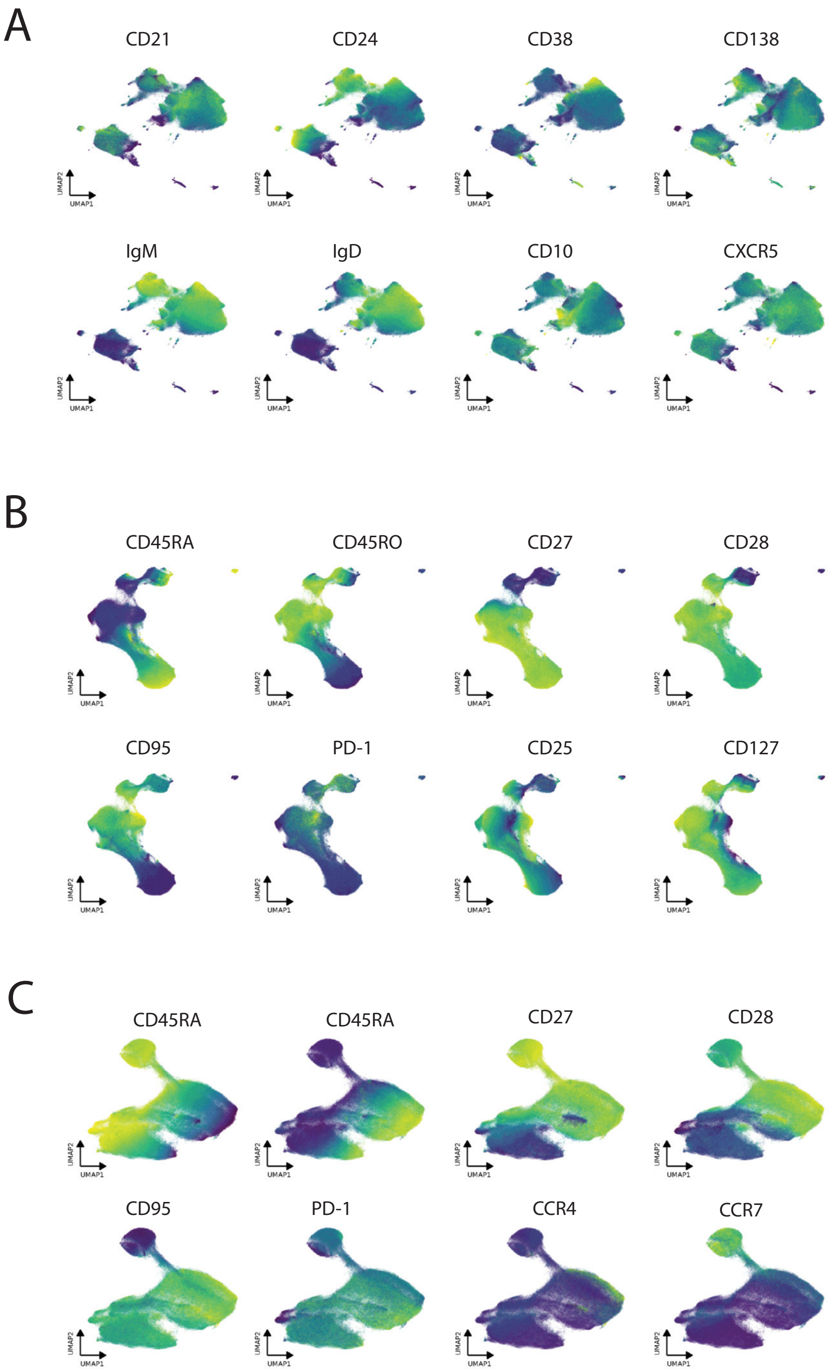

### Figure S7

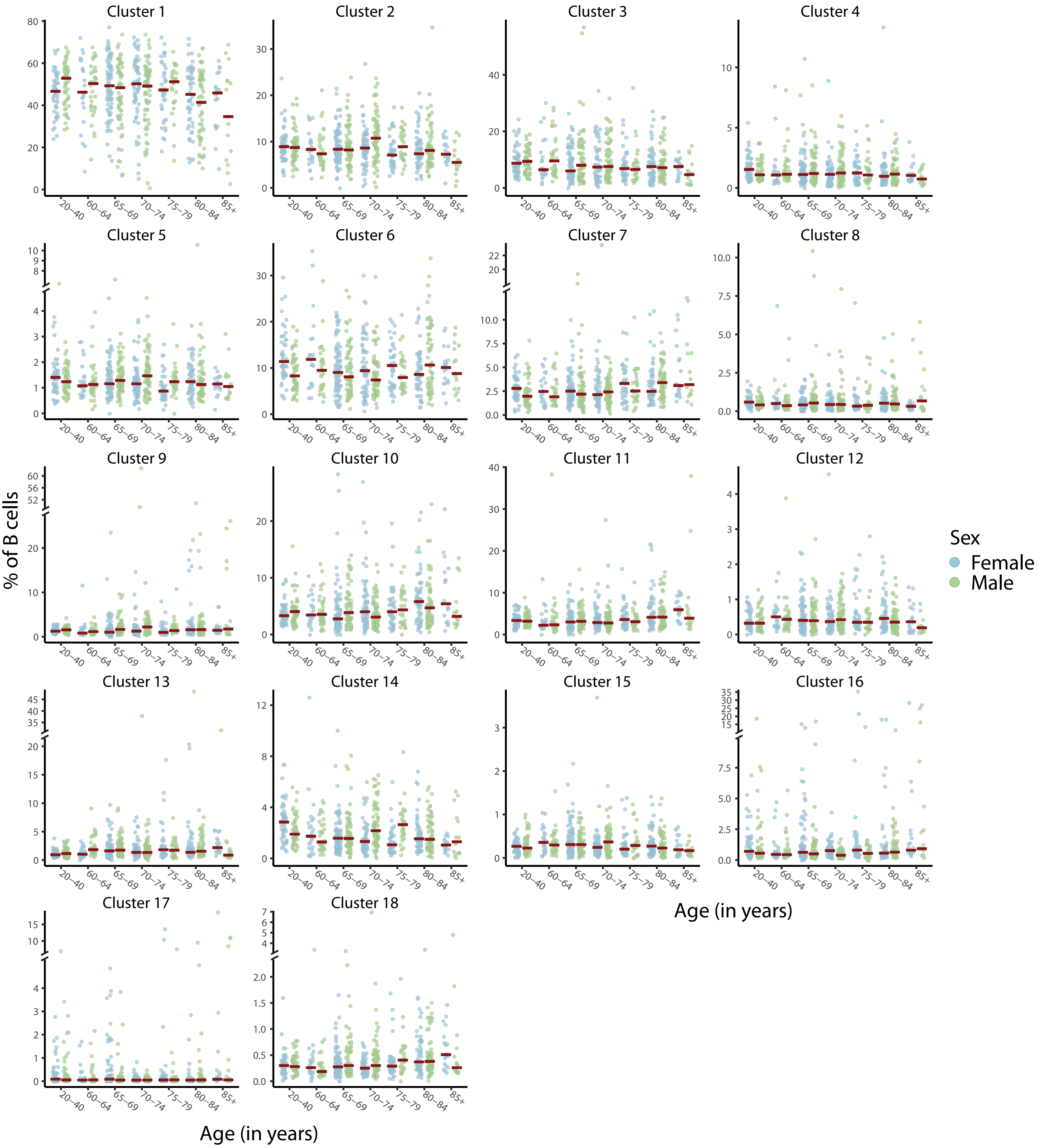

### Figure S8

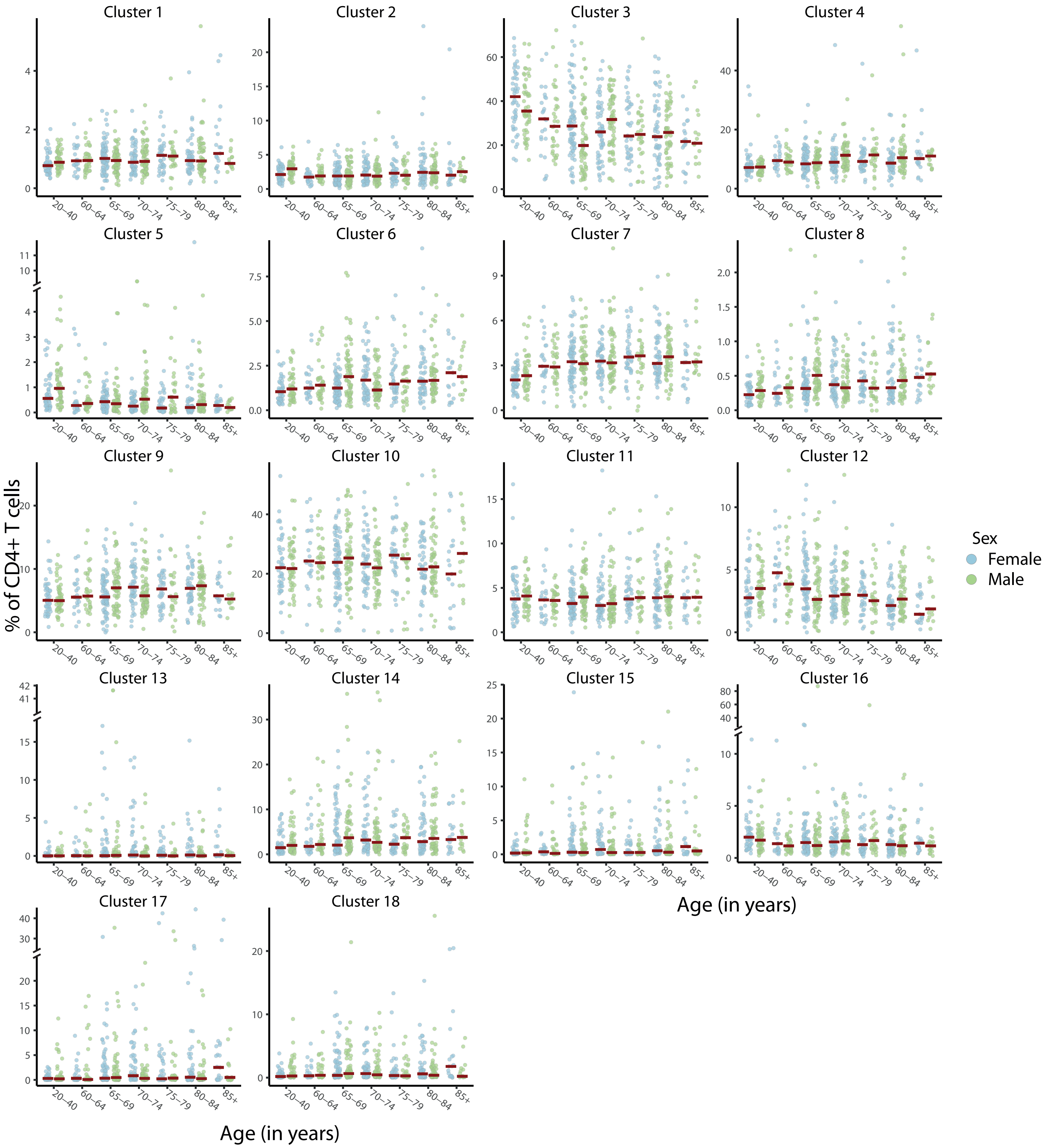

### Figure S9

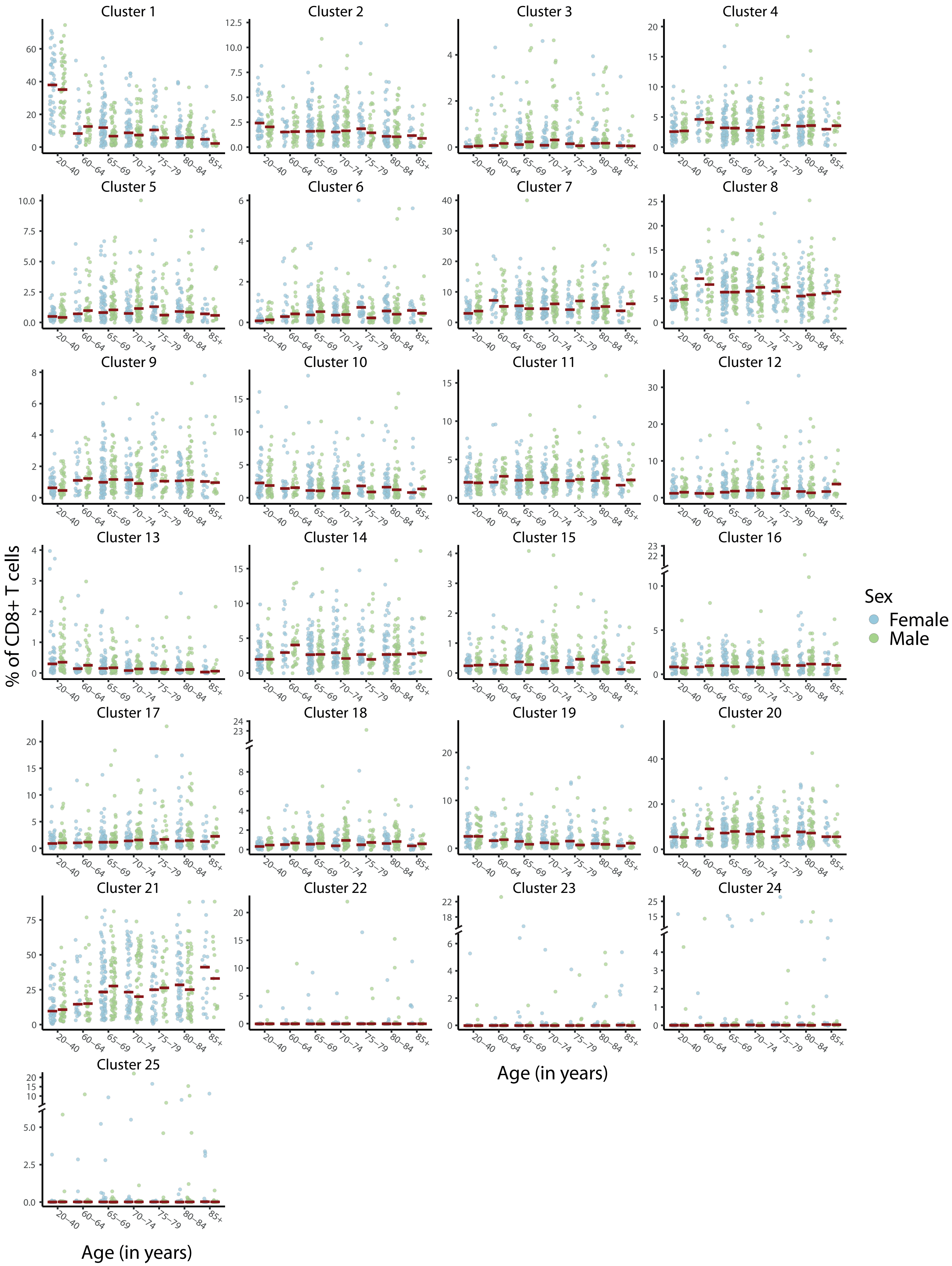

### Figure S10

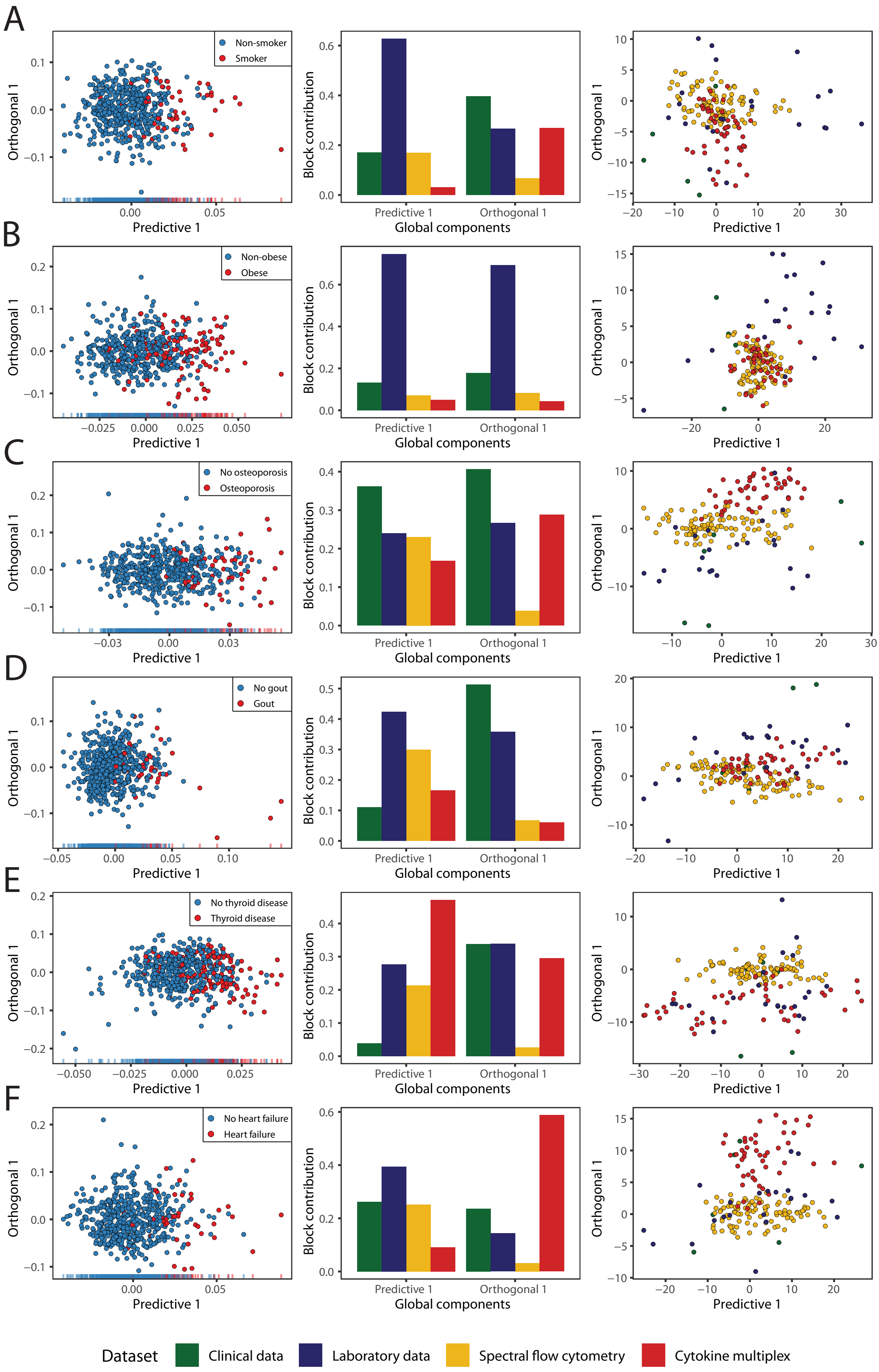
