## Supplementary Tables for "Integrative deep immune profiling of the elderly reveals systems-level signatures of aging, sex, smoking, and clinical traits"

**Suppl. Table 1:** Plasma cytokine concentrations measured by multiplex assay in young adults and elderly in the RESIST SI cohort.

| Cytokine | Plasma concentration (pg/mL)^1^ | | Statistic | Effect size | P Value^2^ |
| --- | --- | --- | --- | --- | --- |
|  | Young adults | Elderly |  |  |  |
| CCL2 | 5.9 (4.2-8.1) | 6.5 (4.9-8.9) | 30482 | 0.08 | 0.034 |
| CCL3 | 0.8 (0.7-0.9) | 0.9 (0.8-1.2) | 38049 | 0.26 | <0.001 |
| CCL4 | 45.2 (39.4-55.5) | 50.5 (42.6-60.6) | 31661.5 | 0.11 | 0.005 |
| CCL5 | 1512.9 (985-2096.3) | 1613.5 (1106-2189.1) | 28700 | 0.04 | 0.283 |
| CCL7 | 0.1 (0-0.2) | 0.1 (0-0.2) | 30078.5 | 0.08 | 0.051 |
| CCL11 | 18.5 (14.1-25) | 23.4 (17-32.1) | 34474 | 0.18 | <0.001 |
| CCL27 | 154.7 (110.1-215.6) | 252.7 (182.1-348.6) | 41218 | 0.33 | <0.001 |
| CXCL1 | 25.6 (25.6-25.6) | 25.6 (25.6-25.6) | 26860.5 | 0 | 0.976 |
| CXCL8 | 0.8 (0.5-1.5) | 1.2 (0.8-1.9) | 34226.5 | 0.17 | <0.001 |
| CXCL9 | 39.5 (32.7-51.2) | 89.1 (63.4-150.7) | 48456.5 | 0.5 | <0.001 |
| CXCL10 | 81.3 (56.8-108.1) | 111.8 (76.8-156.6) | 37519.5 | 0.25 | <0.001 |
| CXCL12 | 245.6 (211.3-300.6) | 235.4 (188.4-279) | 23488 | 0.08 | 0.046 |
| IFN-α2 | 1.7 (1.1-3) | 1.4 (0.6-3) | 26136.5 | 0.02 | 0.654 |
| IFN-**γ** | 2.7 (1.5-4.2) | 3 (1.5-4.7) | 28183.5 | 0.03 | 0.442 |
| IL-1a | 13.9 (11.6-16.9) | 13.9 (10.8-18) | 25862 | 0.02 | 0.549 |
| IL-1b | 0.1 (0-0.2) | 0.2 (0.1-0.2) | 33199.5 | <0.001 | 0.15 |
| IL-1RA | 54.4 (40.3-71) | 61.4 (42.6-85.4) | 30745.5 | 0.09 | 0.023 |
| IL-2 | 0.1 (0.1-0.3) | 0.1 (0.1-0.4) | 28973 | 0.05 | 0.178 |
| IL-2Rα | 29.2 (19.6-38.4) | 29.8 (20.4-43.6) | 29294 | 0.06 | 0.155 |
| IL-3 | 0 (0-0.1) | 0 (0-0.1) | 27682 | 0.02 | 0.563 |
| IL-4 | 1.1 (0.7-1.5) | 1.2 (0.8-1.6) | 30150.5 | 0.08 | 0.054 |
| IL-5 | 20.9 (18.9-20.9) | 20.9 (7.8-20.9) | 26718 | 0 | 0.914 |
| IL-6 | 0.1 (0.1-0.1) | 0.1 (0.1-0.1) | 24903.5 | 0.05 | 0.241 |
| IL-7 | 0.7 (0.7-26.7) | 0.7 (0.7-28.9) | 28205 | 0.539 | 0.03 |
| IL-9 | 103.2 (89.1-119) | 115.3 (94.5-135.3) | 31714 | 0.11 | 0.004 |
| IL-10 | 1.2 (0.6-2.1) | 1.3 (0.7-2.1) | 28660 | 0.04 | 0.294 |
| IL-12(p40) | 5.2 (2-18.2) | 3.7 (2-18.2) | 23700.5 | 0.08 | 0.05 |
| IL-12(p70) | 0 (0-0.2) | 0 (0-0.2) | 26049.5 | 0.02 | 0.549 |
| IL-13 | 0.6 (0.1-1.6) | 0.8 (0.1-1.8) | 28892.5 | 0.05 | 0.233 |
| IL-15 | 46.4 (46.4-46.4) | 46.4 (46.4-46.4) | 26954 | 0.01 | 0.818 |
| IL-16 | 20.3 (16.6-28.4) | 24.1 (18.1-30.3) | 31120.5 | 0.1 | 0.012 |
| IL-17 | 1 (0.6-1.5) | 1.2 (0.7-2) | 30925 | 0.09 | 0.017 |
| IL-18 | 8 (4.6-10.5) | 7.9 (3.9-13.8) | 28199 | 0.03 | 0.539 |
| FGF-β | 0.9 (0.9-2) | 0.9 (0.9-5.4) | 28725 | 0.05 | 0.231 |
| GCSF | 16.4 (11.3-21.3) | 23.3 (16-34.8) | 37679.5 | 0.25 | <0.001 |
| GMCSF | 2.3 (0.1-3.3) | 1.9 (0.2-3.2) | 26076 | 0.02 | 0.692 |
| HGF | 114.9 (93.4-145.2) | 156.1 (116.7-199) | 38257 | 0.26 | <0.001 |
| LIF | 0.8 (0.4-0.8) | 0.8 (0.8-3.2) | 29935.5 | 0.08 | 0.055 |
| MCSF | 3.5 (1.8-5.2) | 4.5 (2.3-8.1) | 32891 | 0.14 | <0.001 |
| MIF | 205 (145.2-287.9) | 187.2 (134.9-260) | 24184 | 0.06 | 0.112 |
| PDGF-bb | 52.6 (39.1-74.2) | 61.9 (41.2-89.7) | 29715 | 0.07 | 0.095 |
| SCF | 13.3 (9.4-16.2) | 14.4 (10.7-19.6) | 31997 | 0.12 | 0.003 |
| SCGF-β | 24978 (18159.2-34298.3) | 23465.5 (17270.2-32639.1) | 25478.5 | 0.03 | 0.41 |
| βNGF | 0 (0-0.1) | 0 (0-0.1)  v | 24109.5 | 0.07 | 0.084 |
| TNFα | 20 (15.9-29) | 21.8 (15.2-29.6) | 28163.5 | 0.03 | 0.449 |
| TNF-β | 81.7 (70-102.3) | 93.3 (75.8-115.6) | 31739.5 | 0.11 | 0.004 |
| TRAIL | 6.7 (5.3-10.4) | 8.4 (5.6-11.7) | 30534 | 0.09 | 0.031 |
| VEGF | 22.7 (22.7-22.7) | 22.7 (22.7-22.7) | 26563 | 0.03 | 0.447 |

^1^ Median and interquartile ranges are displayed.

^2^ Wilcoxon rank sum test

**Suppl. Table 2:** Marker characteristics, frequencies, and variability of HLA-DR^neg^ innate cell clusters.

| Cluster | Subset type | Marker characteristics | Mono-tonic^1^ | Mean % of HLA-DR^neg^ innate cells | | Coefficient of variation^2^ | |
| --- | --- | --- | --- | --- | --- | --- | --- |
|  |  |  |  | Young adults | Elderly | Young adults | Elderly |
| 1 | Early NK | CD16lowCD56highCD127+NKg2D+CD11clow | Yes | 4.3 | 3.4 | 0.74 | 0.79 |
| 2 | Early NK | CD16-CD56^high^CD127+NKg2D-CD11c^low^ | Yes | 0.5 | 0.4 | 1.58 | 1.38 |
| 3 | Early NK | CD16-CD56+CD127-NKG2D+CD11c- | Yes | 2.3 | 2.4 | 0.88 | 1.06 |
| 4 | Mature NK | CD16+CD56+NKG2D+CD11c^low^ | Yes | 45.8 | 45.2 | 0.31 | 0.31 |
| 5 | Mature NK | CD16+CD56+NKG2D+CD11c- | Yes | 38.1 | 39.7 | 0.38 | 0.35 |
| 6 | Mature NK | CD16+CD56+NKG2D+CD80+CD206+ | Yes | 0.6 | 0.4 | 1.83 | 1.97 |
| 7 | Terminal NK | CD16+CD56-NKG2D+ | Yes | 0.8 | 1.2 | 0.90 | 2.66 |
| 8 | CD56−CD16− NK | CD16-CD56-NKG2D+ | No | 0.4 | 0.4 | 0.68 | 2.1 |
| 9 | Basophils | Lin-CD123+CRTH2+CD11c^low^ | No | 3.2 | 3.5 | 1.66 | 1.61 |
| 10 | Mast cells | Lin-CD123^low^CD11c^low^ | No | 0.2 | 0.4 | 1.04 | 1.46 |
| 11 | ILC 1 | Lin-CD127+CRTH2-NKG2D-CD103+ | No | 0.2 | 0.2 | 2.20 | 1.34 |
| 12 | ILC 2 | Lin-CD127+CRTH2+ | No | 0.4 | 0.2 | 0.73 | 0.97 |
| 13 | ILC 3 | Lin-CD127+CRTH2-NKG2D+ | No | 1.3 | 0.9 | 0.48 | 1.18 |
| 14 | ILC 3 | Lin-CD127+CRTH2-NKG2D- | No | 1.7 | 1.6 | 0.57 | 1.98 |
| 15 | Other | CD127+CD206+CD80^high^CD86^low^ | No | 0.1 | 0.1 | 1.93 | 3.16 |

Abbr: ILC: Innate lymphoid cell; NK: Natural killer cell

^1^ Jonckheere-Terpstra test p ≥ 0.05.

^2^ Standard deviation devided by the mean

**Suppl. Table 3:** Marker characteristics, frequencies, and variability of HLA-DR^pos^ innate cell clusters.

| Cluster | Subset type | Marker characteristics | Mono-tonic^1^ | Mean % of HLA-DR^pos^ innate cells | | Coefficient of variation^2^ | |
| --- | --- | --- | --- | --- | --- | --- | --- |
|  |  |  |  | Young adults | Elderly | Young adults | Elderly |
| 1 | Classical Monocyte | CD16-CD14+CD11c+ | Yes | 60.7 | 63.1 | 0.33 | 0.27 |
| 2 | Classical Monocyte | CD16-CD14+CD115+CD103- | No | 3.2 | 2.8 | 1.35 | 1.52 |
| 3 | Classical Monocyte | CD16-CD14+CD115+HLADR+CD103+ | No | 2.3 | 3.4 | 1.20 | 1.14 |
| 4 | Classical Monocytes | CD16^low^CD14+CD103-CCR5-CCR2+ | Yes | 0.6 | 0.8 | 1.16 | 2.02 |
| 5 | Intermediate Monocytes | CD16+CD14^low^CD86+ | No | 1.4 | 2.3 | 0.92 | 0.7 |
| 6 | Intermediate Monocytes | CD16+CD14+CD86+CD11c^high^ | No | 0.4 | 0.8 | 1.20 | 1.2 |
| 7 | Non-classical monocyte | CD16+CD14- | No | 5.2 | 8.1 | 0.77 | 0.74 |
| 8 | Non-classical Monocyte | CD16^low^CD14- | No | 0.7 | 1.1 | 1.16 | 1.23 |
| 9 | Granulo-monocytic precursors | CD16^low^CD14- CD123^low^CD11c^low^ | No | 0.1 | 0.2 | 0.92 | 1.83 |
| 10 | pDC | Lin-CD123+CCR2+CCR5^low^ | No | 14.3 | 8.5 | 1.20 | 1.21 |
| 11 | cDC1 | Lin-CCR2^low^CD11c+CD141+ | Yes | 0.5 | 0.3 | 0.77 | 1.64 |
| 12 | cDC2 | Lin-CCR2+CD11c^high^CCR5+CD1c+ | No | 9.0 | 6.9 | 1.16 | 0.93 |
| 13 | cDC2 | Lin-CCR2+CD11c^high^CCR5+CD1c+CD141+ | Yes | 1.1 | 0.8 | 1.42 | 1.07 |
| 14 | Other | Negative to all measured markers | No | 0.4 | 0.9 | 1.02 | 2.21 |

Abbr: cDC: Conventional dendritic cell; pDC: Plasmacytoid dendritic cell

^1^ Jonckheere-Terpstra test p ≥ 0.05.

^2^ Standard deviation devided by the mean

**Suppl. Table 4:** Marker characteristics, frequencies, and variability of γδ T cell clusters.

| Cluster | Subset type | Marker characteristics | Mono-tonic^1^ | Mean % of γδ T cells | | Coefficient of variation^2^ | |
| --- | --- | --- | --- | --- | --- | --- | --- |
|  |  |  |  | Young adults | Elderly | Young adults | Elderly |
| 1 | Vγ9- Naïve | Vγ9-CD45RA+CD62L^high^CD16- | No | 0.1 | 0.9 | 2.04 | 2.55 |
| 2 | Vγ9- T_EM_ | Vγ9-CD45RA+CD62LlowCD16^mid^ | No | 11.2 | 22.0 | 1.50 | 0.99 |
| 3 | Vγ9+ Naïve | Vγ9+CD45RA+CD27+CD28+/- CD127+ | No | 35.5 | 25.9 | 0.42 | 0.59 |
| 4 | Vγ9+ T_CM_ | Vγ9+CD45RA-CD45RO+CD27+ | No | 33.7 | 31.7 | 0.50 | 0.61 |
| 5 | Vγ9+ T_EM_ | Vγ9+CD45RA-CD45RO+CD27- | Yes | 3.5 | 4.2 | 0.93 | 1.13 |
| 6 | Vγ9+ T_EMRA_ | Vγ9+CD45RA+CD27-CD16+CD56^mid^ | Yes | 15.7 | 15.0 | 0.95 | 1.11 |
| 7 | Vγ9 Other | Vγ9+ markers | Yes | 0.2 | 0.3 | 0.72 | 1.38 |

^1^ Jonckheere-Terpstra test p ≥ 0.05.

^2^ Standard deviation devided by the mean

**Suppl. Table 5:** Marker characteristics, frequencies, and variability of B cell clusters.

| Cluster | Subset type | Marker characteristics | Mono-tonic^1^ | Mean % of B cells | | Coefficient of variation^2^ | |
| --- | --- | --- | --- | --- | --- | --- | --- |
|  |  |  |  | Young adults | Elderly | Young adults | Elderly |
| 1 | B cell Naïve | IgM+IgD+CD27^low^CCR6+CD21+CD24- | No | 48.0 | 45.5 | 0.21 | 0.33 |
| 2 | B cell Naïve | IgM^low^IgD+CD27-CCR6+CD21-CD24- | Yes | 9.2 | 9.0 | 0.40 | 0.49 |
| 3 | Non-switched memory B cells | IgM+IgD^low^CD27+CD21^low^CD24+ | No | 9.8 | 9.1 | 0.47 | 0.75 |
| 4 | Non-switched memory B cells | IgM+IgD+CD27+CD21-CD24+ | No | 1.6 | 1.5 | 0.59 | 0.92 |
| 5 | Non-switched memory B cells | IgM-IgD+CD27^low^CD21-CD24- | Yes | 1.5 | 1.4 | 0.57 | 0.61 |
| 6 | Switched memory B cells | IgM-IgD- CD27+CD21-CD24+CD138+ | Yes | 10.8 | 10.2 | 0.48 | 0.56 |
| 7 | Switched memory B cells | IgM-IgD-CD27^low^CD21-CD24-CD138+ | No | 2.6 | 3.1 | 0.54 | 0.74 |
| 8 | Switched memory B cells | IgM-IgD-CD27+ CD21^low^CD138+ | Yes | 0.6 | 0.7 | 0.62 | 1.42 |
| 9 | Switched memory B cells | IgM-IgD-CD27+ CCR7+CD21-CD24^low^ | No | 1.5 | 2.7 | 0.43 | 1.91 |
| 10 | Switched memory B cells | IgM-IgD-CD27+ CD21+CD24+ | No | 4.1 | 5.0 | 0.58 | 0.83 |
| 11 | Switched memory B cells | IgM-IgD-CD27+CD21-CD24^low^CD138- | No | 3.6 | 4.3 | 0.45 | 0.90 |
| 12 | Transitional B cells | IgM+IgD^low^CD27^low^CD21-CD24+CD38+ | Yes | 0.4 | 0.5 | 0.68 | 0.92 |
| 13 | Transitional B cells | IgM+IgD+CD27^low^CD21-CD24- | No | 1.3 | 2.4 | 0.60 | 1.46 |
| 14 | Transitional B cells | IgM+IgD+CD27^low^CD21+CD24-CXCR5^low^CD138+ | No | 2.5 | 2.0 | 0.56 | 0.78 |
| 15 | Transitional B cells | IgM+IgD+CD27-CD21-CD24+CXCR5^low^CD38+ | Yes | 0.3 | 0.4 | 0.69 | 0.93 |
| 16 | Plasmablasts | CD20-CD38+CD138+CD27+ | Yes | 1.5 | 1.5 | 1.57 | 2.28 |
| 17 | Plasmablasts | CD20-CD38+CD138+CD27- | Yes | 0.4 | 0.4 | 2.09 | 3.85 |
| 18 | Double negative B cells | IgM-IgD-CD27-CD21- | No | 0.3 | 0.4 | 0.66 | 1.18 |

^1^ Jonckheere-Terpstra test p ≥ 0.05.

^2^ Standard deviation devided by the mean

**Suppl. Table 6:** Marker characteristics, frequencies, and variability of CD4+ T cell clusters.

| Cluster | Subset type | Marker characteristics | Mono-tonic^1^ | Mean % of CD4+ T cells | | Coefficient of variation^2^ | |
| --- | --- | --- | --- | --- | --- | --- | --- |
|  |  |  |  | Young adults | Elderly | Young adults | Elderly |
| 1 | CD4+ Naïve | CD45RA+CCR7+CD25+CD127- | No | 0.9 | 1.1 | 0.37 | 0.55 |
| 2 | CD4+ Naïve | CD45RA+CCR7+CD25+CD127+ | Yes | 2.7 | 2.4 | 0.42 | 0.80 |
| 3 | CD4+ Naïve | CD45RA+CCR7+CD127+ | No | 39.4 | 27.6 | 0.34 | 0.52 |
| 4 | CD4+ Naïve | CD45RA+CCR7+CD45RO^mid^CD127+ | No | 8.0 | 10.4 | 0.57 | 0.53 |
| 5 | CD4+ Naïve | CD45RA+CCR7+CD25-CD127- | No | 1.0 | 0.6 | 0.88 | 1.56 |
| 6 | CD4+ Tregs | CD45RA-CD25+CD27+CD127-CD62L+ | No | 1.2 | 1.8 | 0.57 | 0.70 |
| 7 | CD4+ Tregs | CD45RA-CD25+CD27+CD127+CD62L- | No | 2.3 | 3.5 | 0.45 | 0.44 |
| 8 | CD4+ Tregs | CD45RA-CD25+CD27-CD127-CD62L+ | No | 0.3 | 0.5 | 0.59 | 0.74 |
| 9 | CD4+ T_CM_ | CD45RA-CCR7+CD27+CD28+  CCR4-CCR6-CD127-CXCR3^mid^ | No | 5.5 | 6.9 | 0.46 | 0.49 |
| 10 | CD4+ T_CM_ | CD45RA-CCR7+CD27+CD28+  CCR4-CCR6-CD127+CXCR3^mid^ | Yes | 23.0 | 24.0 | 0.39 | 0.42 |
| 11 | CD4+ T_CM_ | CD45RA-CCR7+CD27-CCR4+  CCR6+ | Yes | 4.3 | 4.2 | 0.56 | 0.59 |
| 12 | CD4+ Tfh-like | CD45RA-CCR7+CXCR5+ | No | 3.5 | 3.3 | 0.50 | 0.61 |
| 13 | CD4+ iNKTs | CD45RO+CCR7-CD56+ | No | 0.2 | 0.9 | 2.67 | 3.09 |
| 14 | CD4+ T_EM_ | CD45RO+CD95+CD127+ | No | 2.9 | 4.9 | 1.10 | 1.13 |
| 15 | CD4+ T_EM_ | CD45RO+CD95-CD127- | No | 0.8 | 1.6 | 2.07 | 1.88 |
| 16 | CD4+ T_EMRA_ | CD45RA+CCR7^low^CD27+CD28+ | No | 2.3 | 2.2 | 0.75 | 2.29 |
| 17 | CD4+ T_EMRA_ | CD45RA+CCR7-CXCR3-CD27-CD28- | No | 0.9 | 2.7 | 2.00 | 2.17 |
| 18 | CD4+ T_EMRA_ | CD45RA+CD45RO+CCR7+CD27-CD28- | No | 0.8 | 1.6 | 1.72 | 1.76 |

^1^ Jonckheere-Terpstra test p ≥ 0.05.

^2^ Standard deviation devided by the mean

**Suppl. Table 7:** Marker characteristics, frequencies, and variability of CD8+ T cell clusters.

| Cluster | Subset type | Marker characteristics | Mono-tonic^1^ | Mean % of CD8+ T cells | | Coefficient of variation^2^ | |
| --- | --- | --- | --- | --- | --- | --- | --- |
|  |  |  |  | Young adults | Elderly | Young adults | Elderly |
| 1 | CD8+ Naïve | CD45RA+CCR7+CD28^mid^CD95- | No | 36.3 | 10.9 | 0.49 | 0.95 |
| 2 | CD8+ Naïve | CD45RA+CCR7+CD28+CD95^low^ | No | 2.5 | 2.0 | 0.64 | 0.86 |
| 3 | CD8+ T_CM_ | CD45RA^low^+CCR7+CD28^high^CCR4+CD95+ | No | 0.2 | 0.5 | 1.93 | 1.67 |
| 4 | CD8+ T_CM_ | CD45RA^low^CD45RO+CCR7-PD1+ | Yes | 3.1 | 3.9 | 0.60 | 0.66 |
| 5 | CD8+ T_CM_ | CD45RA^low^CD45RO^mid^CCR7^low^CD25+ | No | 0.7 | 1.5 | 0.97 | 1.02 |
| 6 | CD8+ T_CM_ | CD45RA-CD45RO+CCR7^mid^CCR4+CD25+ | No | 0.2 | 0.6 | 1.28 | 1.17 |
| 7 | CD8+ T_CM_ | CD45RA-CD45RO+CCR7-PD1+ | Yes | 4.5 | 6.4 | 0.74 | 0.73 |
| 8 | CD8+ T_EM_ | CD45RA^low^CD45RO^low^CCR7- | Yes | 5.2 | 7.1 | 0.44 | 0.56 |
| 9 | CD8+ T_EM_ | CD45RA-CD45RO+CCR7^low^CD25+ | No | 0.8 | 1.5 | 0.87 | 0.83 |
| 10 | CD8+ T_EM_ | CD45RA^low^CD45RO+CD127^high^CCR6+ | No | 3.0 | 2.0 | 0.96 | 1.18 |
| 11 | CD8+ T_EM_ | CD45RA^low^CD45RO+CD27^low^CD28^low^ | Yes | 2.4 | 2.8 | 0.65 | 0.66 |
| 12 | CD8+ T_EM_ | CD45RA^low^CD45RO+CD27-CD28-CD127- | No | 2.1 | 3.1 | 0.95 | 1.22 |
| 13 | CD8+ T_EM_ | CD45RA^low^CD45RO^low^CD27-CD28+CD127^high^CCR6+Tim3+ | No | 0.6 | 0.3 | 1.26 | 1.47 |
| 14 | CD8+ T_EM_ | CD45RA-CD45RO+PD1+ | No | 2.4 | 3.4 | 0.78 | 0.80 |
| 15 | CD8+ T_EM_ | CD45RA-CD45RO+CD27^low^CD28+ | Yes | 0.3 | 0.4 | 0.83 | 1.09 |
| 16 | CD8+ T_EM_ | CD45RA-CD45RO+CD27^mid^CD28+ | No | 1.1 | 1.3 | 0.91 | 1.14 |
| 17 | CD8+ T_EM_ | CD54RA^low^CD45RO+CD27-CD28-CD127- | No | 1.7 | 2.3 | 1.14 | 1.26 |
| 18 | CD8+ T_EM_ | CD45RA-CD45RO+CD27-CD28+ | No | 0.5 | 1.0 | 0.92 | 1.39 |
| 19 | CD8+ T_EMRA_ | CD45RA^low^CCR7-CD27+CD28+CD127^high^CCR6+Tim3+ | No | 3.3 | 1.9 | 0.88 | 1.26 |
| 20 | CD8+ T_EMRA_ | CD45RA+CCR7-CD27+CD28-CD127^low^ | No | 5.9 | 8.6 | 0.65 | 0.73 |
| 21 | CD8+ T_EMRA_ | CD45RA+CCR7-CD27-CD28-CD127- | No | 15.1 | 28.7 | 0.84 | 0.70 |
| 22 | CD8+ T_EMRA_ | CD45RA+CCR7-CD27+CD28+CD127+ | Yes | 7.7 | 9.2 | 0.42 | 0.47 |
| 23 | CD8+ T_EMRA_ | CD45RA+CCR7-CD27-CD28-CD127^low^ | No | 0.1 | 0.2 | 6.98 | 6.92 |
| 24 | CD8+ T_EMRA_ | CD45RA+CD45RO^mid^CCR7-CD27+CD28+CD127+ | No | 0.2 | 0.4 | 7.00 | 5.93 |
| 25 | CD8+ T_EMRA_ | CD45RA+CD45RO+CCR7-CD27+CD28+CD127+ | Yes | 0.1 | 0.3 | 5.60 | 5.56 |

^1^ Jonckheere-Terpstra test p ≥ 0.05.

^2^ Standard deviation devided by the mean

**Suppl. Table 8:** Innate immune cell antibody panel.

| **Specificity** | **Fluorochrome** | **Clone** | **Company** | **Cat#** | **Dilution** |
| --- | --- | --- | --- | --- | --- |
| CD45 | BUV395 | HI30 | BD | 563792 | 1:200 |
| CD16 | BUV496 | 3G8 | BD | 612944 | 1:100 |
| CD14 | BUV563 | MoP9 | BD | 741441 | 1:100 |
| CD115 | BUV615 | 94D2 | BD | 751279 | 1:50 |
| CD11c | BUV661 | B-ly6 | BD | 612967 | 1:100 |
| CD56 | BUV737 | NCAM16.2 | BD | 612766 | 1:100 |
| CD86 | BUV805 | BU63 | BD | 748375 | 1:50 |
| CD123 | Super Bright 436 | 6H6 | Invitrogen | 62-1239-42 | 1:50 |
| CD103 | BV480 | Ber-ACT8 | BD | 746472 | 1:50 |
| CCR5 | BV510 | J418F1 | BioLegend | 359128 | 1:50 |
| HLA-DR | BV570 | L243 | BioLegend | 307638 | 1:50 |
| CD141 | BV605 | M80 | BioLegend | 344114 | 1:20 |
| CD80 | BV650 | 2D10 | BioLegend | 305227 | 1:100 |
| CD125 | BV786 | A14 | BD | 743932 | 1:100 |
| CD1c | AF488 | L161 | BioLegend | 331522 | 1:20 |
| CD3 | AF532 | UCHT1 | Invitrogen | 58-0038-42 | 1:50 |
| CD127 | PerCP-Cy5.5 | HIL-7R-M21 | BD | 560551 | 1:100 |
| NKg2D | PE | 1D11 | BioLegend | 320805 | 1:100 |
| CTRH2 | PE-Dazzle 594 | BM16 | BioLegend | 350125 | 1:50 |
| CD19 | PE-Cy7 | SJ25C1 | BD | 557835 | 1:200 |
| CCR2 | APC | K036C2 | BioLegend | 357208 | 1:100 |
| CD66b | AF647 | G10F5 | BioLegend | 305110 | 1:100 |
| CD64 | AF700 | 10.1 | BD | 561188 | 1:50 |
| CD206 | APC Fire750 | 15-2 | BioLegend | 321134 | 1:50 |
| Viability | Zombie NIR |  | BioLegend | 423106 | 1:400 |

**Suppl. Table 9:** Adaptive immune cell antibody panel.

| **Specificity** | **Fluorochrome** | **Clone** | **Company** | **Cat#** | **Dilution** |
| --- | --- | --- | --- | --- | --- |
| CD45RA | BUV395 | HI100 | BD | 740298 | 1:200 |
| CD16 | BUV496 | 3G8 | BD | 612944 | 1:100 |
| CD4 | BUV563 | RPA-T4 | BD | 741353 | 1:200 |
| CCR4 | BUV615 | 1G1 | BD | 613000 | 1:20 |
| CD21 | BUV661 | 1048 | BD | 750187 | 1:100 |
| CD56 | BUV737 | NCAM16.2 | BD | 612766 | 1:100 |
| CD27 | BUV805 | L128 | BD | 748704 | 1:100 |
| CD28 | BV421 | 2H7 | BioLegend | 302330 | 1:100 |
| CD45RO | Pacific Blue | UCHL1 | BioLegend | 304216 | 1:100 |
| IgD | BV480 | IA6-2 | BD | 566138 | 1:200 |
| CD138 | BV510 | 145/15 | Miltenyi | 130-113-623 | 1:50 |
| CD20 | BV570 | L243 | BioLegend | 307638 | 1:100 |
| TCRVg9 | BV605 | B3 | BD | 744036 | 1:100 |
| CXCR3 | BV650 | G025H7 | BioLegend | 353730 | 1:100 |
| CCR6 | BV711 | G034E3 | BioLegend | 353436 | 1:100 |
| CXCR5 | BV750 | RF8B2 | BD | 747111 | 1:200 |
| CCR7 | BV785 | G043H7 | BioLegend | 353230 | 1:50 |
| PD-1 | FITC | NAT105 | BioLegend | 367412 | 1:20 |
| CD3 | AF532 | UCHT1 | Invitrogen | 58-0038-42 | 1:50 |
| CD8 | SparkBlue550 | SK1 | BioLegend | 344760 | 1:200 |
| CD19 | PerCP | HIB19 | BioLegend | 302228 | 1:20 |
| CD14 | BB700 | MφP9 | BD | 566465 | 1:100 |
| CD62L | PerCP-Cy5.5 | DREG56 | BioLegend | 304824 | 1:100 |
| CD38 | PerCP-eF710 | HB7 | Invitrogen | 46-0388-42 | 1:100 |
| CD10 | PE | HI10a | BioLegend | 312204 | 1:100 |
| CD24 | PE-Dazzle 594 | ML5 | BioLegend | 311134 | 1:100 |
| CD95 | PE-Cy5 | DX2 | Biolegend | 305610 | 1:20 |
| CD25 | Biotin/SaV-PE-Cy5.5 | BC96 | BioLegend | 302624 | 1:100 |
| CTLA4 | PE-Cy7 | L3D10 | BioLegend | 349914 | 1:20 |
| HLA-DR | PE-Fire810 | L243 | BioLegend | 307683 | 1:50 |
| TCRgd | APC | 11F2 | Miltenyi | 130-113-500 | 1:50 |
| IgM | AF647 | MHM-88 | BioLegend | 314536 | 1:100 |
| CD127 | APC R700 | A019D5 | BioLegend | 351344 | 1:100 |
| Viability | Zombie NIR |  | BioLegend | 423106 | 1:400 |
| Tim3 | APC Fire750 | F38-2E2 | BioLegend | 345044 | 1:100 |
